# Gatekeeper helix activates Golgi SM protein Sly1 and directly mediates close-range vesicle tethering

**DOI:** 10.1101/2020.01.16.906719

**Authors:** M. Duan, R.L. Plemel, T. Takenaka, A. Lin, B.M. Delgado, U. Nattermann, D.P. Nickerson, J. Mima, E.A. Miller, A.J. Merz

## Abstract

The essential Golgi protein Sly1 is a member of the SM (<u>S</u>ec1/<u>m</u>ammalian Unc-18) family of SNARE chaperones. Sly1 was originally identified through gain-of-function alleles that bypass requirements for diverse vesicle tethering factors. Employing genetic analyses and chemically defined reconstitutions of ER-Golgi fusion, we discovered that a loop conserved among Sly1 family members is not only autoinhibitory, but also acts as a positive effector. An amphipathic helix within the loop directly binds high-curvature membranes; membrane binding is required for relief of Sly1 autoinhibition and allows Sly1 to directly tether incoming vesicles to the Qa-SNARE on the target organelle. The *SLY1-20* allele bypasses requirements for diverse tethering factors but loses this functionality if Sly1 membrane binding is impaired. We propose that long-range tethers, including Golgins and multisubunit tethering complexes, hand off vesicles to Sly1, which then tethers at close range to activate SNARE assembly and fusion in the early secretory pathway.

## INTRODUCTION

Traffic through the secretory and endocytic systems depends on accurate and timely targeting of transport vesicles to acceptor organelles. The terminal stage of targeting is membrane fusion, catalyzed by the formation of *trans*-SNARE complexes that zipper together, doing the mechanical work of moving two membranes into proximity and driving their merger. Although SNAREs alone can drive fusion and confer some compartmental selectivity, spontaneous SNARE assembly is slow and error-prone. Consequently, an array of tethering factors and SNARE chaperones are indispensable *in vivo* (Baker and Hughson, 2016; Gillingham and Munro, 2019). For example, every SNARE-mediated fusion event that has been closely examined requires a cofactor of the Sec1/mammalian Unc-18 (SM) family.

For decades the mechanisms of SM protein function were enigmatic (Carr and Rizo, 2010; Rizo and Sudhof, 2012; Sudhof and Rothman, 2009) but biochemical work, structural studies, and single-molecule force spectroscopy suggest that SM proteins are assembly chaperones for *trans*-SNARE complex formation, and that SMs act, at least in part, by templating the initial SNARE zippering reaction (Baker et al., 2015; Jiao et al., 2018) and by protecting appropriately formed prefusion complexes from kinetic proofreading by the SNARE disassembly proteins Sec17/α-SNAP and Sec18/NSF (Lobingier et al., 2014; Ma et al., 2013; Schwartz et al., 2017; Xu et al., 2010). There are four subfamilies of SM proteins. The budding yeast *Saccharomyces cerevisiae* has one representative of each. Vps33, the first SM identified genetically, controls fusion at late endosomes and lysosomes (Banta et al., 1990; Patterson, 1932; Sevrioukov et al., 1999). Vps45 controls fusion at early endosomal compartments (Cowles et al., 1994; Piper et al., 1994). Sec1 and its orthologs (Unc-18/Munc-18) control exocytosis (Grote et al., 2000; Novick et al., 1979; Verhage et al., 2000; Wu et al., 1998). Finally, fusion at the Golgi, and probably at the endoplasmic reticulum (ER), is controlled by Sly1 (Li et al., 2005; Lupashin et al., 1996; Ossig et al., 1991; Peng and Gallwitz, 2002; Sogaard et al., 1994).

The genetics of *SLY1* are complex and revealing. Ypt1 (yeast Rab1) is an essential regulator of docking and fusion at the Golgi. *SLY1* was originally identified through an allele, *SLY1-20*, that dominantly <u>S</u>uppresses the <u>L</u>ethality of <u>Y</u>pt1 deficiency (Dascher et al., 1991; Ossig et al., 1991; Ossig et al., 1995). Subsequent work by several groups showed that *SLY1-20* suppresses deficiencies not only of Ypt1, but of numerous other factors that promote ER and Golgi traffic. These include the Dsl complex (*dsl1* was originally identified through its genetic interaction with *SLY1-20*; Reilly et al., 2001; Vanrheenen et al., 2001); the COG complex (*cog2, cog3*; VanRheenen et al., 1998; VanRheenen et al., 1999); the TRAPP complexes (*bet3-1*; Sacher et al., 1998); the Golgin coiled-coil tether Uso1 (yeast p115; Sapperstein et al., 1996); Ypt6 (yeast Rab6) and its nucleotide exchange complex (*ric1*; Bensen et al., 2001; Li et al., 2007); and the Ypt6 effector complex GARP (*vps53*; Vanrheenen et al., 2001). In addition, *SLY1-20* suppresses partial deficiencies of Golgi SNAREs (*sec22*; Ossig et al., 1991); COPI coat subunits *sec21*; (Ossig et al., 1991); and the COPI Arf GAP Glo3 (Vanrheenen et al., 2001).

*SLY1-20* and the similar allele *SLY1-15* encode missense substitutions at adjacent positions within a loop insertion that is evolutionarily conserved among Sly1 subfamily members, but absent from the other three SM subfamilies (Dascher et al., 1991; Li et al., 2007). On this basis it was hypothesized that the Sly1 loop is auto-inhibitory, and that *SLY1-20* and related alleles gain function by releasing the loop from its closed, autoinhibitory state (Bracher and Weissenhorn, 2002; Li et al., 2007). This proposal was strengthened by the discovery that the Sly1 loop occludes a conserved site which, in the lysosomal SM Vps33, binds R/v-SNAREs with high affinity (Baker et al., 2015).

The physiological mechanism by which the Sly1 loop’s putative auto-inhibitory activity is released to promote SNARE complex formation is unknown, but was suggested to require Ypt1, the yeast Rab1 ortholog (Bracher and Weissenhorn, 2002; Li et al., 2007). Here, we show that the loop’s inhibitory activity is released when an amphipathic helix within the loop interacts directly with the incoming vesicle membrane’s lipid bilayer. Moreover, the loop allows Sly1 to directly tether incoming vesicles: the Sly1 N-lobe is anchored to the Qa/t-SNARE on the target organelle, while the Sly1 regulatory loop binds the vesicle lipid bilayer. We propose that this membrane binding steers Sly1 into an orientation optimal for productive R/v-SNARE association and *trans*-complex assembly. This schema explains how Sly1-20 can bypass the otherwise essential functions of so many different Golgi tethering factors, and suggests that the Sly1 regulatory loop links Sly1 activation, the capture of transport vesicles addressed to organelles of the early secretory pathway, and productive *trans-*SNARE complex assembly.

## RESULTS

### New *SLY1* alleles define a regulatory loop in Sly1

We thought it likely that early screens which identified *SLY1\** bypass alleles were not saturated, and that a more focused screen might yield additional informative alleles. Uso1 is a Golgin-class tether that is a direct effector of Ypt1/Rab1. Loss of Uso1 is lethal, and this lethality is suppressed by *SLY1-20* (Ballew et al., 2005; Sapperstein et al., 1996). We therefore designed a selection for dominant *SLY1\** alleles that could suppress the loss of *USO1* (**Fig. 1A**). (In this report, sets of *SLY1* alleles and their products are referred to collectively as *SLY1\** and Sly1*.) Our screen retrieved many *SLY1\** alleles, most carrying multiple missense substitutions. From these, individual missense substitutions were re-introduced into wild type *SLY1* and tested for their ability to suppress deficiencies of Uso1 or Ypt1 (**Fig. 1B**; **Supplementary Table 1**). Importantly, our screen retrieved the original *SLY1-20* and *SLY1-15* alleles. We also identified suppressing substitutions at nearby sites on helix α20, and on the short segment linking helices α20 and α21. Additionally, we identified suppressing substitutions at the base of the Sly1-specific loop, and at positions cradling the base of the loop, but non-adjacent within the linear polypeptide sequence. One of these was T559I. A genomic survey for gene pairs exhibiting spontaneous suppressing interactions found that a substitution at the same position, T559K, dominantly suppressed deficiencies of both the GARP subunit Vps53 and the Arf GAP Glo3 (van Leeuwen et al., 2016). Most of the gain-of-function single substitutions that we tested suppressed *ypt1-3* but, in contrast to the multi-site mutants obtained in the initial selection for *uso1Δ* bypass, were unable to suppress *uso1Δ*. (**Supplementary Fig. S1; Supplementary Table 1**). Thus, strong Sly1 gain-of-function phenotypes can arise through individual substitutions or through the compounded effects of multiple weak driver substitutions. These results show that earlier screens were not, as had been suggested, saturated (Li et al., 2007).

**Fig. 1.**
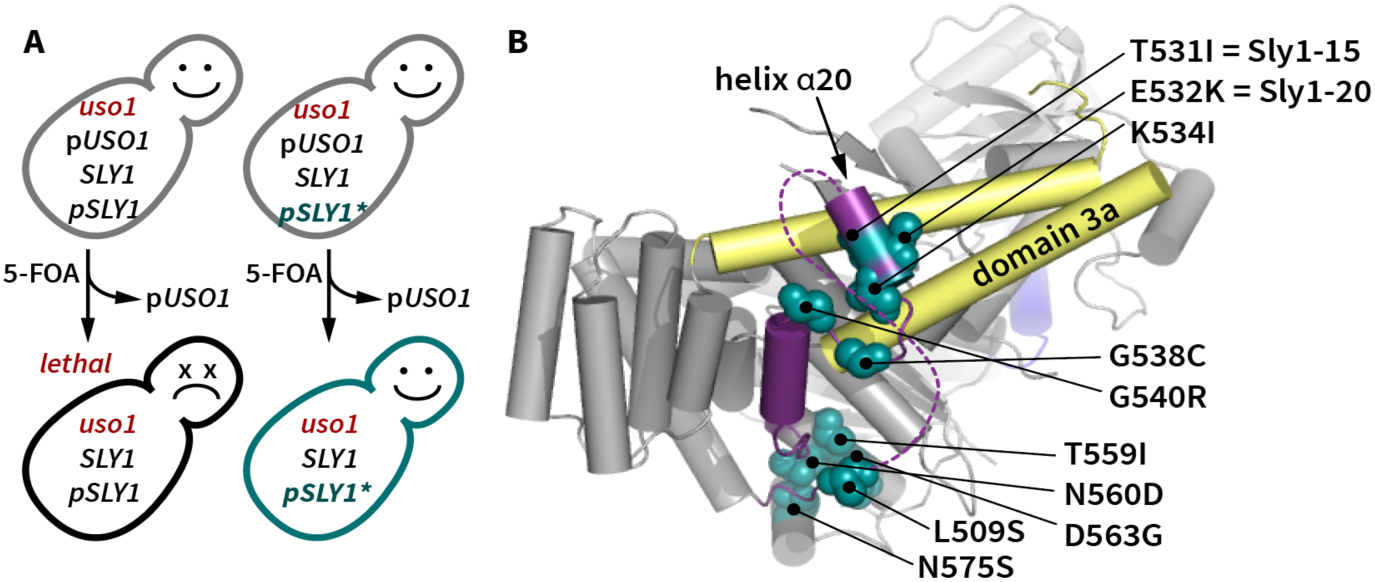
New gain-of-function *SLY1* alleles. **A.** Selection used in this study. A library of *SLY1\** alleles was constructed by PCR-based mutagenesis and cloned into a single-copy plasmid. The library was then transformed into a *SLY1 uso1*Δ strain, with *USO1* provided on a balancer plasmid bearing the counterselectable *URA3* marker. Ejection of p*USO1* was forced by 5-fluoroorotic acid (5-FOA). This positively selects viable cells carrying dominant mutant *SLY1*\* alleles that bypass the otherwise essential *USO1* requirement. **B.** Locations within Sly1 (PDB: 1MQS) of single missense substitutions that suppress requirements for Ypt1, or for both Ypt1 and Uso1. The loop is indicated in purple, with the dashed line denoting the portion of the loop not resolved in the structure. Yellow shading indicates the domain 3a helical hairpin which, by analogy to Vps33 and Munc18-1, is hypothesized to scaffold assembly of Qa- and R-SNARE trans-complexes.

As noted by Baker and coworkers (2015), helices α20 and α21 sit atop two conserved regions that in Vps33 are of special importance for SNARE binding: domain 3a, which serves as a scaffold to nucleate the parallel, in-register assembly of the Qa- and R-SNAREs, and an aromatic pocket that serves as a high-affinity anchoring point for the R-SNARE juxtamembrane linker. On the basis of the Vps33 structures and the original *SLY1-20* and *SLY1-15* alleles, Baker *et al*. (2015) speculated that when closed, the Sly1 loop might prevent R-SNARE binding to Sly1. The dominant suppressors obtained in our screen and data presented below reinforce and extend that model.

### Sly1 bypass suppressors are hyperactive in a minimal fusion system

*In vivo* genetic tests and crude *in vitro* transport systems (Baker et al., 1988; Ballew et al., 2005; Ruohola et al., 1988) cannot tell us whether Sly1* mutants must interact with additional proteins beyond the core SNARE fusion machinery to manifest gain of function. To overcome this limitation, we developed a chemically defined reconstituted proteoliposome (RPL) system to monitor fusion driven by ER-Golgi SNAREs (**Fig. 2A**). This system, adapted from an assay developed to study homotypic vacuole fusion (Zucchi and Zick, 2011), employs two orthogonal pairs of Förster resonance energy transfer (FRET) probes, to simultaneously monitor both lipid and content mixing in small (20 µL) reaction volumes. Although we present only content mixing data in this manuscript, the lipid mixing signal provides an intrinsic control, allowing us to detect partial hemifuison or fusion that is accompanied by lysis.

**Fig. 2:**
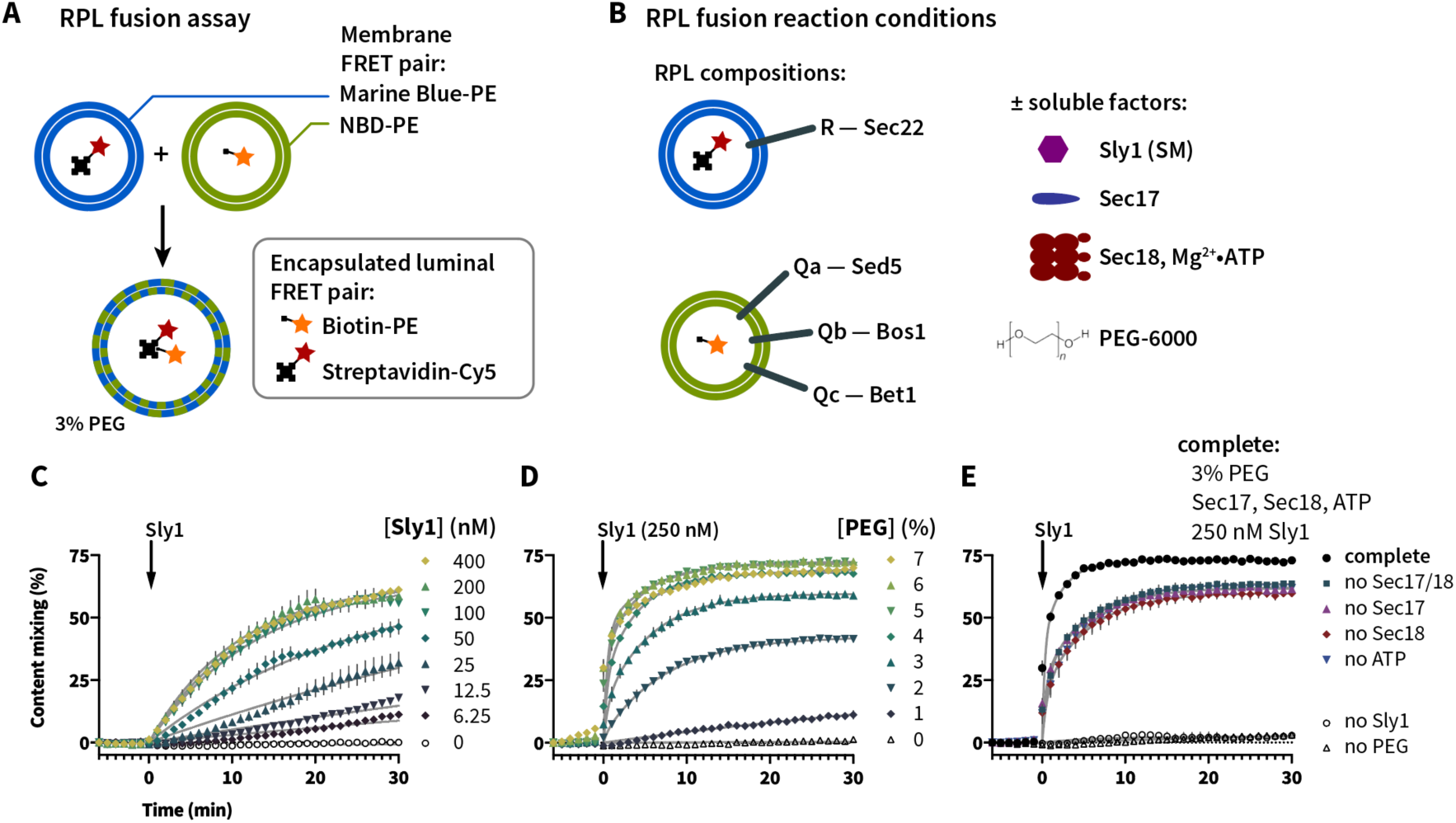
Setup and characterization of the *in vitro* fusion system. **A**. Reporter systems for lipid and content mixing. RPLs are prepared with encapsulated content mixing FRET pair, and with the membranes doped with an orthogonal FRET pair. **B**. SNARE topology of the RPLs used in this study, and soluble components added to stimulate fusion. **C-E**, characterization of the system using content mixing readout. **C**, Requirement for Sly1. Reactions were set up with Q- and R-SNARE RPLs, and 3% PEG. Fusion activity was monitored for 5 min., and then Sly1 was added at time = 0 (arrows) to the indicated final concentrations. Note that fusion activity is saturated at 100 nM [Sly1]. **D**. Requirement for tethering. Reactions were set up with Q- and R-SNARE RPLs, with the indicated final concentrations of PEG. Fusion activity was monitored for 5 min., and then Sly1 was added to a final concentration of 250 nM. Note that at 6% and 7% PEG, some Sly1-independent fusion occurs prior Sly1 addition. **E**. Effects of the SNARE disassembly machinery. Reactions were set up with Q- and R-SNARE RPLs, and with or without PEG, Sec17, Sec18, ATP, and Sly1, as indicated. Fusion was initiated by adding Sly1. For **C-E**, points show mean ± s.e.m. of three independent experiments; in many cases the error bars are smaller than the symbols. Gray lines show least-squares nonlinear fits of a second-order kinetic model.

In previous work SMs were shown to stimulate SNARE-mediated lipid mixing, but only in the presence of tethering factors or molecular crowding agents that substitute for tethering factors (Furukawa and Mima, 2014; Yu et al., 2015). Consistent with these previous studies, content mixing in heterotypic reactions between RPLs bearing the R-SNARE Sec22, and RPLs bearing the Q-SNAREs Sed5, Bos1 and Bet1, was strongly stimulated only when both Sly1 and a crowding agent (polyethylene glycol 6000; PEG) were provided (**Fig. 2C,D**). Under these experimental conditions the stimulatory effect of Sly1 saturated at 100 nM. Two other studies have reported *in vitro* stimulation of fusion by Sly1, though at a 45× higher concentration of Sly1 than the 100 nM used in most of our experiments (Furukawa and Mima, 2014; Jun and Wickner, 2019). Pre-incubation of the RPLs with Mg^2+^·ATP and the SNARE disassembly chaperones Sec17 and Sec18 (yeast α-SNAP and NSF) resulted in immediate and almost complete fusion upon Sly1 addition (**Fig. 2E**), likely indicating that SNAREs on the RPLs equilibrate between productive and refractory configurations, and that Sec17/18-mediated disassembly shifts this equilibrium toward productive, Sly1-reactive configurations.

Next, we compared the activity of wild type Sly1 to three bypass suppressors: Sly1-20 and two of the new alleles identified in our screen. Each variant was tested in reactions containing 3% or 0% polyethylene glycol-6000 (PEG). At 3% PEG all four Sly1 variants drove fusion with similar efficiency (**Fig. 3A**). In marked contrast, at 0% PEG (**Fig. 3B**) all three Sly1 suppressor mutants drove fusion substantially more efficiently than the wild type. In reactions containing Sec17, Sec18, and Mg^2+^·ATP (**Fig. 3C,D**) the same overall pattern emerged. As PEG bypasses requirements for tethering factors and potentiates SM-mediated fusion, our results show for the first time that these Sly1 gain-of-function mutants are intrinsically hyperactive, requiring only SNAREs (or SNAREs and disassembly chaperones) to stimulate fusion, and not additional cellular factors such as Rabs or tethering factors. These results directly mirror the *in vivo* genetic suppression patterns observed for *SLY1-*20 and otherwise essential vesicle tethering regulators and effectors.

**Fig. 3:**
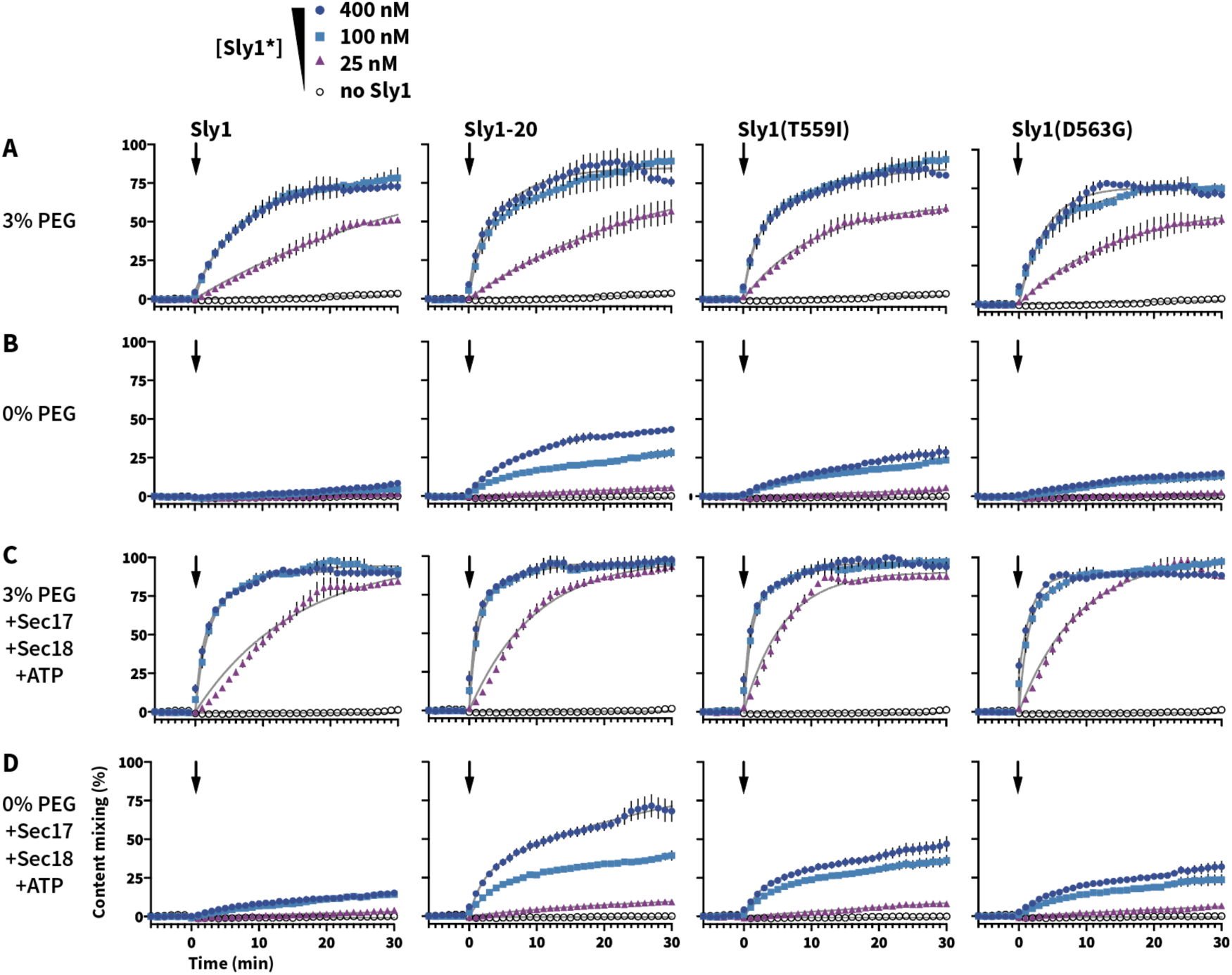
Gain-of-function Sly1 mutants alleviate the tethering requirement *in vitro*. Reactions were set up as in Fig. 2, with the initial mixture containing R-SNARE and Qabc-SNARE RPLs and, as indicated for each row of panels, 0 or 3% PEG, and in the absence or presence of Sec17, Sec18 (both 100 nM), and ATP (1 mM). After a 5 min incubation, wild type Sly1 or the indicated mutants were added (arrows) at 0, 25, 100, or 400 nM to initiate fusion. Points show the mean ± s.e.m. from three or more independent experiments; in many cases the error bars are smaller than the symbols. Gray lines show least-squares nonlinear fits of a second-order kinetic model.

### The Sly1 regulatory loop has positive as well as negative functions

If the Sly1 loop is autoinhibitory, we might predict that excision of the entire loop should hyperactivate Sly1 as much as or more than the suppressing mutations characterized above. To test this hypothesis, we used the *ROSETTA* software environment (Leaver-Fay et al., 2011) to design a set of Sly1 variants in which the loop is replaced by short peptide linkers (**Fig. 4A; Supplementary Table 2**). Surprisingly, each of the “loopless” *sly1* mutants tested *in vivo* exhibited recessive lethality or slow growth in the presence of the counterselection agent 5-FOA at 30° C; temperature sensitivity; and an inability to bypass deficiencies in *YPT1* or *USO1*. One mutant, *sly1-0_2*, exhibited somewhat more robust growth compared to the other alleles when present as the sole copy of *SLY1*. *sly1-0_2* was therefore called *sly1Δloop* and subjected to further scrutiny.

**Fig. 4.**
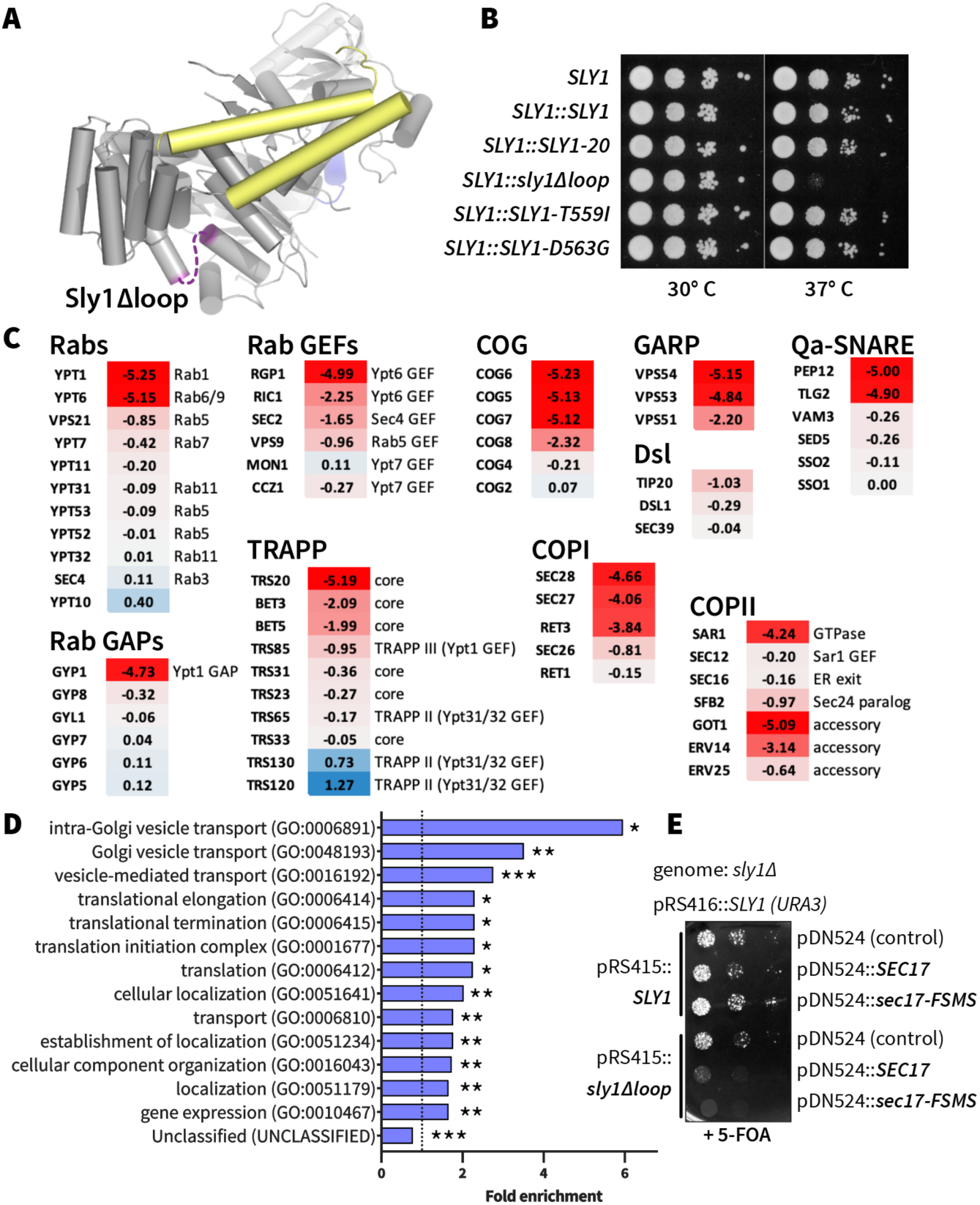
The Sly1 regulatory loop has a positive function *in vivo*. **A**, Diagram showing the location of Sly1 loop replacement with short synthetic linker regions (dashed line; model derived from PDB 1MQS). Sequences of the linker insert designs, and growth phenotypes of the corresponding mutants, are presented in Supplementary Table 2. The domain 3a SNARE assembly template is shown in yellow. **B**, The *sly1Δloop* mutant is temperature sensitive for growth. Dilutions of liquid cultures were spotted as 10x serial dilutions onto YPD agar plates and incubated for 2 days at 30° or 37° C. These are knock-ins at the genomic *SLY1* locus, in the Y8205 strain background used for SGA analysis. **C**, Selected SGA results. Genes exhibiting synthetic interactions with *sly1Δloop* are shown. Scores indicate log*_e_* synthetic growth defects (red) or intergenic suppression (blue). A score of –4.6 indicates a 100× synthetic growth defect. Complete SGA results are presented in Supplementary Dataset 1. **D**, Gene Ontology (GO) Overrepresentation Test of the *sly1Δloop* SGA dataset. Genes with log*_e_* synthetic defect scores ≤ –0.5 were included in the analysis. Bars show all GO-Slim Biological Process categories with statistically significant enrichment scores (*p < 0.05; **p < 10^-2^; ***p < 10^-6^). P-values were calculated using Fisher’s exact test and adjusted for multiple comparisons (Bonferroni’s correction; count = 732). Additional details are presented in Supplementary Dataset 1. **E** *SEC17* overproduction is toxic in cells expressing *sly1Δloop*. *sly1Δ* mutant cells were maintained with a counterselectable *SLY1* balancer plasmid and transformed with single-copy plasmids bearing either *SLY1* or *sly1Δloop*, as well as plasmids carrying *SEC17* or *sec17-FSMS* (Schwartz and Merz, 2009). The balancer plasmid was ejected by plating dilutions on media containing 5-FOA and growth was assayed after 2 days of growth at 30° C.

To gain genome-scale insight into the *sly1Δloop* allele’s loss of function, we used synthetic genome array (SGA) analysis. SGA measures the synthetic sickness or rescue (suppression) of a query allele versus a genome-scale collection of loss-of-function alleles (Tong and Boone, 2005). *sly1Δloop* was knocked in at the genomic *SLY1* locus. The resulting strain grew normally on rich YPD medium containing 5-FOA at 30° C, but slowly compared to strains with wild-type or hyperactive *SLY1* alleles at 37° C (**Fig. 4B**). When subjected to SGA analysis, *sly1Δloop* had a synthetic-sick or synthetic-lethal interaction with ten of twelve genes previously reported to exhibit positive suppressing interactions with *SLY1-20*, as well as with dozens of additional genes that function in organelle biogenesis and membrane traffic — especially traffic into and through the Golgi (**Fig. 4C; Supplementary Dataset 1**). Gene Ontology (GO) analysis (Mi et al., 2019; The Gene Ontology Consortium, 2019) recvealed that enrichment for these gene functions was both selective and statistically significant (**Fig. 4D**). Wild-type *SLY1* activity is also required for resistance to the toxic effects of *SEC17* overproduction (Lobingier et al., 2014; Schwartz et al., 2017), and *SEC17* overproduction caused a severe recessive growth defect in *sly1Δloop* cells (**Fig. 4E**). Overexpressino of a mutant, *sec17-FSMS*, caused an even more severe growth defect. Together, the genetic and functional genomic results show that *sly1Δloop* is a recessive loss-of-function allele and not, as expected, a dominant suppressor.

To assess the molecular mechanism of loss-of-function in Sly1Δloop, we returned to the chemically defined fusion system. As shown in **Fig.5**, Sly1Δloop elicited substantially slower fusion compared to wild-type Sly1. Moreover, Sly1Δloop was unable to bypass the tethering requirement *in vitro* (**Figs. 5A and C**), consistent with its inability to suppress Ypt1 and Uso1 deficiencies *in vivo*. Importantly, in dose-response experiments both Sly1Δloop and wild-type Sly1 exhibited saturating fusion activity at ∼100 nM (compare Sly1Δloop in **Fig. 5** to wild type Sly1 in **Fig. 3**). Moreover, Sly1Δloop was properly folded as indicated by circular dichroism spectroscopy (**Fig. 5E**). Thus, the fusion defect of Sly1Δloop is not due to a major fraction of misfolded protein, and the *sly1Δloop* allele is probably not a simple hypomorph. Instead Sly1Δloop is, on a mole-per-mole basis, a less efficient stimulator of SNARE-mediated fusion compared to the wild type. Taken together, the *in vivo* and *in vitro* data argue that the Sly1 regulatory loop is not only autoinhibitory, but that it also must harbor a positive fusion-promoting activity.

**Fig. 5.**
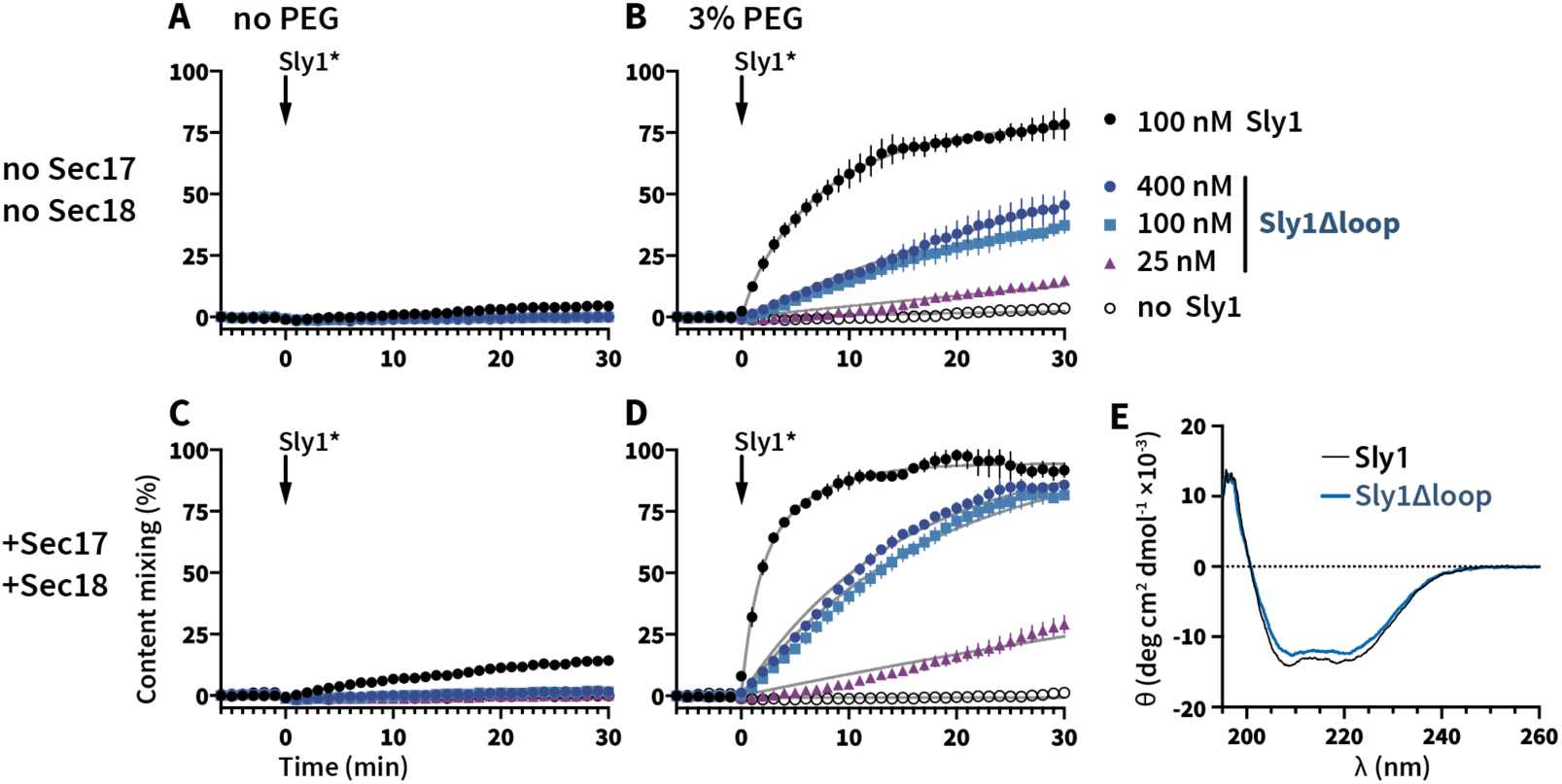
The Sly1 regulatory loop has a positive function *in vitro*. **A-D**, fusion activity of Sly1Δloop versus wild type Sly1. Master mixes were assembled as in Fig. 3, and incubated for 5 min at 30° C. Fusion was initiated by adding (arrows) Sly1 or Sly1Δloop at the concentrations indicated in the legend adjacent to panel **B**. Reactions were run in the absence (**A,C**) or presence (**B,D**) of 3% PEG; and in the absence (**A,B**) or presence (**C,D**) of Sec17, Sec18 (both 100 nM), and ATP. Points show mean ± s.e.m. from three or more independent experiments; in some cases, the error bars are smaller than the symbols. Gray lines show least-squares nonlinear fits of a second-order kinetic model. **E**, Purified Sly1Δloop protein is folded. Circular dichroism spectra of wild type Sly1 and Sly1Δloop. The spectra are normalized to account for small differences in molecular mass and concentration.

### The loop’s positive function resides within ALPS-like helix α21

The regulatory loop’s most highly conserved region is helix α21 (**Fig. 6A, B**). Interestingly, none of the activating gain-of-function mutations isolated to date map to α21. On closer inspection we found that α21 is amphipathic (**Fig. 6C**). We therefore designed a mutant, Sly1-pα21, in which helix α21 is mutated to render it polar rather than amphipathic. Unexpectedly, the *sly1-pα21* allele caused recessive extremely slow growth or lethality (**Fig. 6D**) — a phenotype markedly more severe than that conferred by the *sly1Δloop* allele, which lacks the loop altogether.

**Fig. 6.**
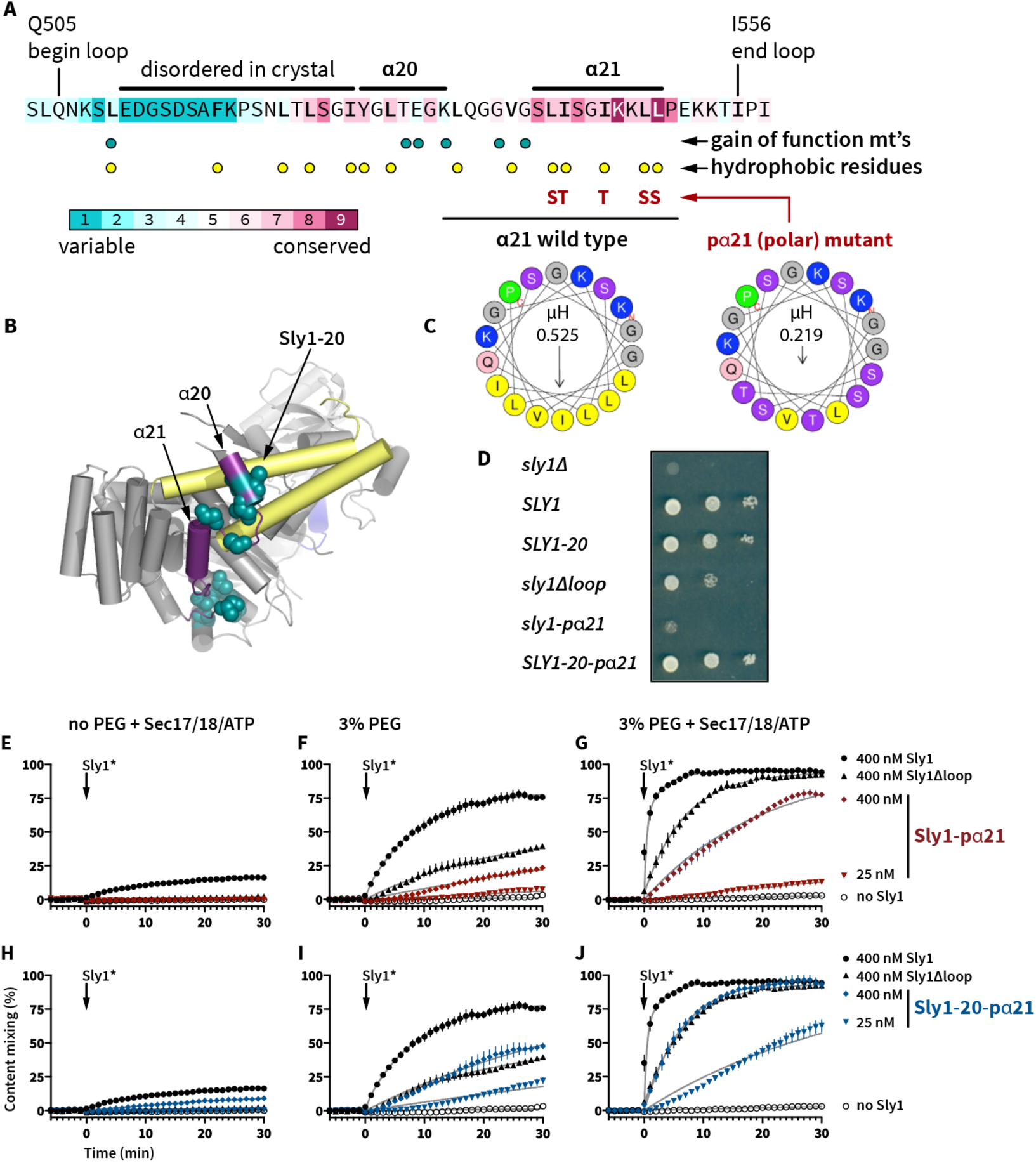
Amphipathic helix α21 is indispensable for normal Sly1 function. **A**, *CONSURF* analysis of evolutionary conservation within the Sly1 loop. Helix α21 is the most highly conserved portion of the loop. Locations of gain-of-function mutations, and hydrophobic residues within the loop, are indicated, as are the five substitutions in the Sly1-pα21 mutant. **B**, Position of helix α21 within Sly1. Note that no gain-of-function mutations within α21 have been identified. The loop is purple; the domain 3a templating domain is yellow. **C**, Helix α21 and residues immediately upstream have the potential to fold into a strongly amphipathic α-helix. The helical wheel renderings comprise the region underlined in black and were produced using *HELIQUEST*; hydrophobic moment (µH) is indicated. **D**, Growth phenotypes of cells carrying *sly1-pα21*, *SLY1-20-pα21*, and other alleles were assayed in a *sly1Δ* strain with a *SLY1* balancer plasmid, which is ejected in the presence of 5-FOA. (**E-J**)RPL fusion with (**E-G**) Sly1-pα and (**H-J**) the compound mutant Sly1-20-pα21. For reference, fusion is also plotted for Sly1 and Sly1^Δloop^. Reactions were set up with (**E,H**) 0% PEG, Sec17 and Sec18 (100 nM each), and ATP (1 mM); (**F,I**) 3% PEG and no Sec17, Sec18 (100 nM each), or ATP; or (**G,J F**) 0% PEG, Sec17 and Sec18 (100 nM each), and ATP (1 mM). Fusion was initiated at time = 0 by adding Sly1 or its mutants, at the concentrations indicated in the legends at the right side of the figure. Points show mean ± s.e.m. from three or more independent experiments; in many cases the error bars are smaller than the symbols. Gray lines show least-squares nonlinear fits of a second-order kinetic model.

Amphipathic helices operate as membrane recognition modules across a wide range of proteins, particularly within the early secretory pathway (Bigay and Antonny, 2012). This suggests a working model: the amphipathic α21 helix probes for the presence of an incoming vesicle and binds to the vesicle’s membrane, pulling the loop away from the R-SNARE binding site. This disinhibits Sly1, allowing Sly1 to catalyze *trans*-SNARE complex assembly. In this model, the Sly1-pα21 protein is nonfunctional because α21 cannot recognize incoming vesicle membranes — and the loop is therefore locked into its auto-inhibited state. To test this hypothesis, we engineered a compound mutant, *SLY1-20-pα21*. This allele comprises both the activating Sly1-20 mutation (E532K) in α20, and the five α21-polar substitutions (**Fig. 6d**). Remarkably, *SLY1-20-pα21* cells exhibited wild type growth (**Fig. 6c**). Unlike *SLY1-20*, however, *SLY1-20-pα21* was unable to suppress the lethality of *ypt1-*3 or *uso1Δ* deficiencies (**Supplementary Table 1**), establishing that the amphipathic character of helix α21 is essential for Sly1-20 hyperactivity *in vivo*.

The *in vivo* results were closely mirrored in fusion experiments with RPLs (**Fig. 6 E-G**). Under every condition tested, Sly1-pα21 was less efficient at stimulating fusion than Sly1Δloop. Fusion in the presence of Sly1-pα21 was reduced in the absence or presence of PEG, as well as in the presence or absence of Sec17, Sec18, and ATP. In contrast to Sly1-pα21, the compound mutant Sly1-20-pα21 (**Fig. 6 H-J**) exhibited a greater ability to stimulate fusion under each of the tested conditions. Importantly, the behaviors of Sly1Δloop and the Sly1-20-pα21 compound mutant were similar. Hence, the amphipathic character of helix α21 is required for the loop’s positive functions: activation and normal function of wild type Sly1, as well as hyperactivity of Sly1-20, both *in vivo* and *in vitro*.

To further test the hypothesis that the regulatory loop has a positive function we prepared chimeras, with fragments of the loop appended to the amino terminus of the Sly1Δloop mutant (**Fig. 7A; Supplementary Table 3**). *In vivo*, chimeras bearing the entire loop, or α20-21, or α21 alone, restored normal growth to Sly1Δloop (**Fig. 7B**). Mutation of five hydrophobic residues within α21 eliminated rescue by *loop-SLY1Δloop* or by *α20-21-SLY1Δloop*. However, at 30°C the polar mutant *pα21-SLY1Δloop* grew almost as well as *α21-SLY1Δloop*. The mechanism of rescue by this mutant construct is unclear.

**Fig. 7.**
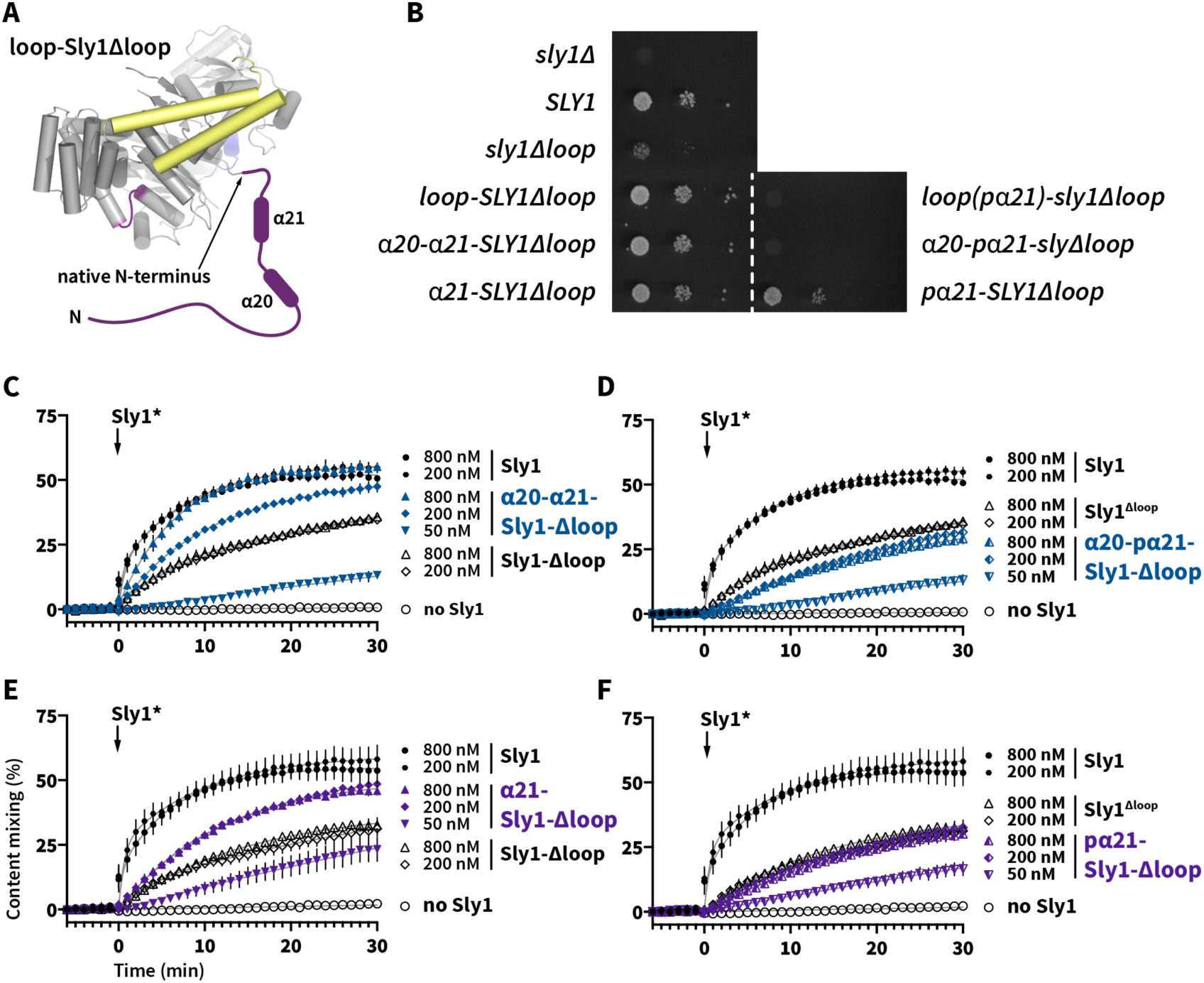
Appending the Sly1 loop to the amino terminus of Sly1Δloop partially restores function. **A**. Chimeric constructs were prepared with different fragments of the Sly1 loop appended to the N-terminus of Sly1 *via* a short, flexible linker (see Supplementary Table 3 for details). Mutants designated pα21 had the five polar substitutions in the appended loop, as described in Fig. 6c. **B**. The loop-Sly1 mutants were expressed from the native *SLY1* promoter on single-copy plasmids. Growth of a *sly1Δ* strain was assessed in the presence of the indicated constructs following ejection of a *SLY1* balancer plasmid by plating on media containing 5-FOA. **C-F**. Fusion driven by mutants with fragments of the loop (C,E) or polar derivatives of these fragments (D,F). Points show mean ±s.e.m. of three independent experiments; in many cases the error bars are smaller than the symbols. Gray lines show least-squares nonlinear fits of a second-order kinetic model.

*In vitro* the α20-21-Sly1Δloop chimera drove almost wild-type fusion when added at 800 nM, whereas its polar mutant (α20-pα21-Sly1Δloop) phenocopied Sly1Δloop. Moreover, in contrast to the result obtained *in vivo*, α21-Sly1Δloop exhibited gain of function relative to Sly1Δloop, while its polar mutant pα21-Sly1Δloop eliminated its gain-of-function relative to Sly1Δloop. Overall (with the interesting exception of the *pα21-SLY1Δloop* allele’s *in vivo* phenotype), these results indicate that that the evolutionarily conserved portion of the Sly1 loop can partially replace the loop’s positive function, even when appended to Sly1 at a non-native location.

### Helix α21 can bind lipid bilayers directly, with preference for high curvature

The above data suggest, but don’t prove, that Sly1 helix α21 binds to membranes, and that the loop allows Sly1 to tether vesicles. To test whether α21 binds to membranes we used a FRET assay. A peptide was synthesized comprising α21 and flanking residues, with an N-terminal tetramethylrhodamine fluorophore (TMR-α21). A control peptide, TMR-pα21, contained the same five substitutions as the Sly1-pα21 mutant (see **Fig. 6A**). Small unilamellar vesicles (SUVs) were prepared with 0.8% Texas Red-phosphatidylethanolamine (TRPE) to serve as a FRET acceptor for TMR. Representative emission spectra for the peptides and SUVs are shown in **Fig. 8A**. SUVs were prepared in two nominal diameters (30 and 200 nm), and with either 6.7% or 30% ergosterol. When mixed with TRPE-doped SUVs, the α21-TMR peptide generated a reproducible FRET signal, evident mainly as donor quenching (**Fig. 8B,C**). Under the same conditions, the control TMR-pα21 peptide exhibited smaller FRET signals. Moreover, the TMR-α21 peptide yielded a larger FRET signal with smaller SUVs, in both the 6.7% and 30% ergosterol conditions (**Fig. 8C**). In contrast, the TMR-pα21 FRET signals did not depend on the SUV diameter. We conclude that helix α21 binds directly to membranes through a mechanism involving the apolar residues within α21, and that it prefers to bind membranes with higher curvature. This is reminiscent of the behavior of ALPS domains, proposed to operate as membrane selectivity filters in the early secretory pathway. However, helix α21 and the Sly1 loop have a higher fraction of charged residues, and the apolar side chains are less bulky, than in canonical ALPS domains (Drin and Antonny, 2010). These differences may explain the apparent insensitivity of TMR-α21 binding to sterol concentration.

**Fig. 8.**
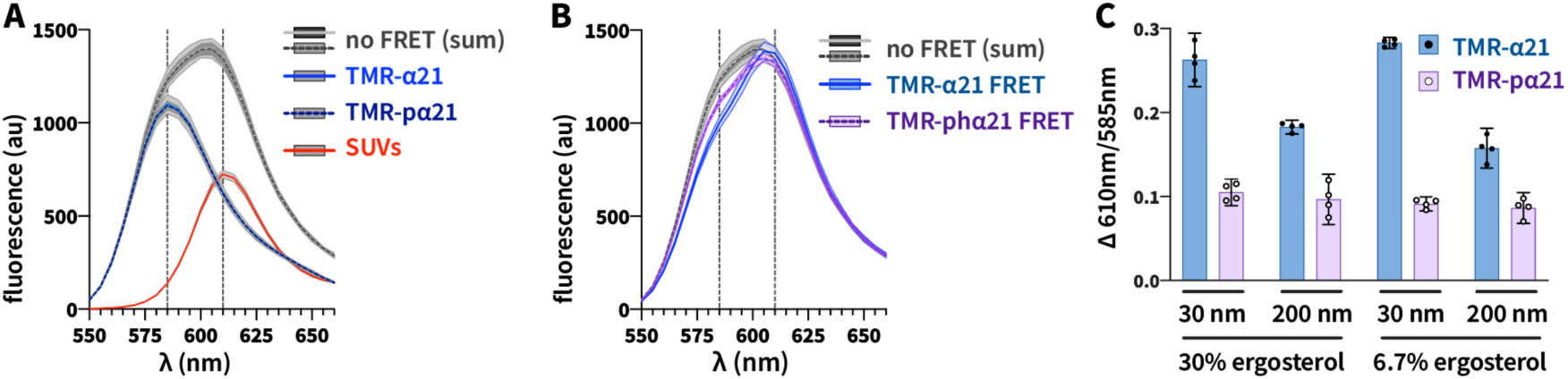
Sly1 helix α21 binds membranes, with a preference for higher curvature. TMR-α21 and TMR-pα21 peptides were added to SUVs of nominal diameter 30 and 200 nm, which contained 1% TRPE as a fluorescence acceptor. **A**. Emission spectra of peptides or liposomes (30 nm diameter, 6.7% ergosterol) measured separately, and the sums of the peptide and liposome spectra. The sums represent the no-FRET condition. Both the TMR-α21 and TMR-pα21 spectra are plotted; they overlap almost exactly. Vertical dashed lines at 585 nm and 610 nm indicate emission peaks for labeled peptides and SUVs, respectively. **B**. Example of FRET data. Spectra from binding reactions containing SUVs (30 nm diameter, 6.7% ergosterol, 500 µM total lipid) and 25 µM TMR-α21 or TMR-pα21 are shown. The no-FRET condition is shown for reference. **C**. Normalized FRET ratios for binding reactions containing the indicated combinations of SUVs and peptides, as in panel B. Traces and bars in A-C show means and ±95% confidence bands from 4 independent experiments.

### Hyperactive Sly1* directly tethers vesicles to the Qa-SNARE

Sly1 binds to the N-peptide-Habc domain of Sed5 (residues 1-210) with sub-nM affinity (Demircioglu et al., 2014; Grabowski and Gallwitz, 1997; Yamaguchi et al., 2002). Thus, we hypothesized that Sly1 may tether heterotypically, with one side of Sly1 binding to the N-peptide of Sed5 on the target membrane while the other side, via helix α21, binds directly to the membrane of the R-SNARE-bearing vesicle. To test this hypothesis we adapted a bead-based assay (**Fig. 9A**) previously used to study Rab-mediated tethering (Lo et al., 2012). First, GST-Sed5 cytoplasmic domain (GST-Sed5_cyt_) or control GST protein were adsorbed to glutathione-agarose beads. Then Sly1* wild type or mutant proteins were added and allowed to bind to the immobilized GST-Sed5_cyt_. Finally, fluorescent vesicles or RPLs were added to the beads, incubated, and imaged by confocal microscopy. If Sly1 or its mutants mediate tethering between Sed5 and the membranes, the signal will be visible in confocal microscopy sections as an equatorial ring of fluorescence on the beads. Qualitative results with wild-type and mutant forms of Sly1 are shown in **Fig. 9A**. To quantify this tethering a bead spin-down assay was used (**Fig. 9B-D**). When Sly1-20, Sly1-T559I, or Sly1-D563G was added to the beads, robust tethering of SNARE-free small unilamellar vesicles (SUVs) was observed (**Figs. 9A,B**). Tethering was eliminated if either Sly1 or Sed5 (“GST control”) was omitted. Tethering was dramatically attenuated with wild type Sly1, with Sly1Δloop, or with Sly1-pα21. An intermediate tethering signal was observed with Sly1-20-pα21. The partial tethering observed with this compound mutant might be due to eight hydrophobic residues on the loop that are still present in the Sly1-pα21 and Sly1-20-pα21 mutants (see **Fig. 6C**). Robust tethering therefore requires that the loop be present, that the loop be open, and that helix α21 be amphipathic. Moeover, as in the peptide binding assays, Sly1-20 mediated tethering was most efficient with small-diameter vesicles, and was insensitive to sterol concentration (compare **Figs. 8C and 9C**). Together these findings indicate that both helix α21 in isolation, and the Sly1 loop in the context of Sly1-20, sense membrane curvature.

**Fig. 9.**
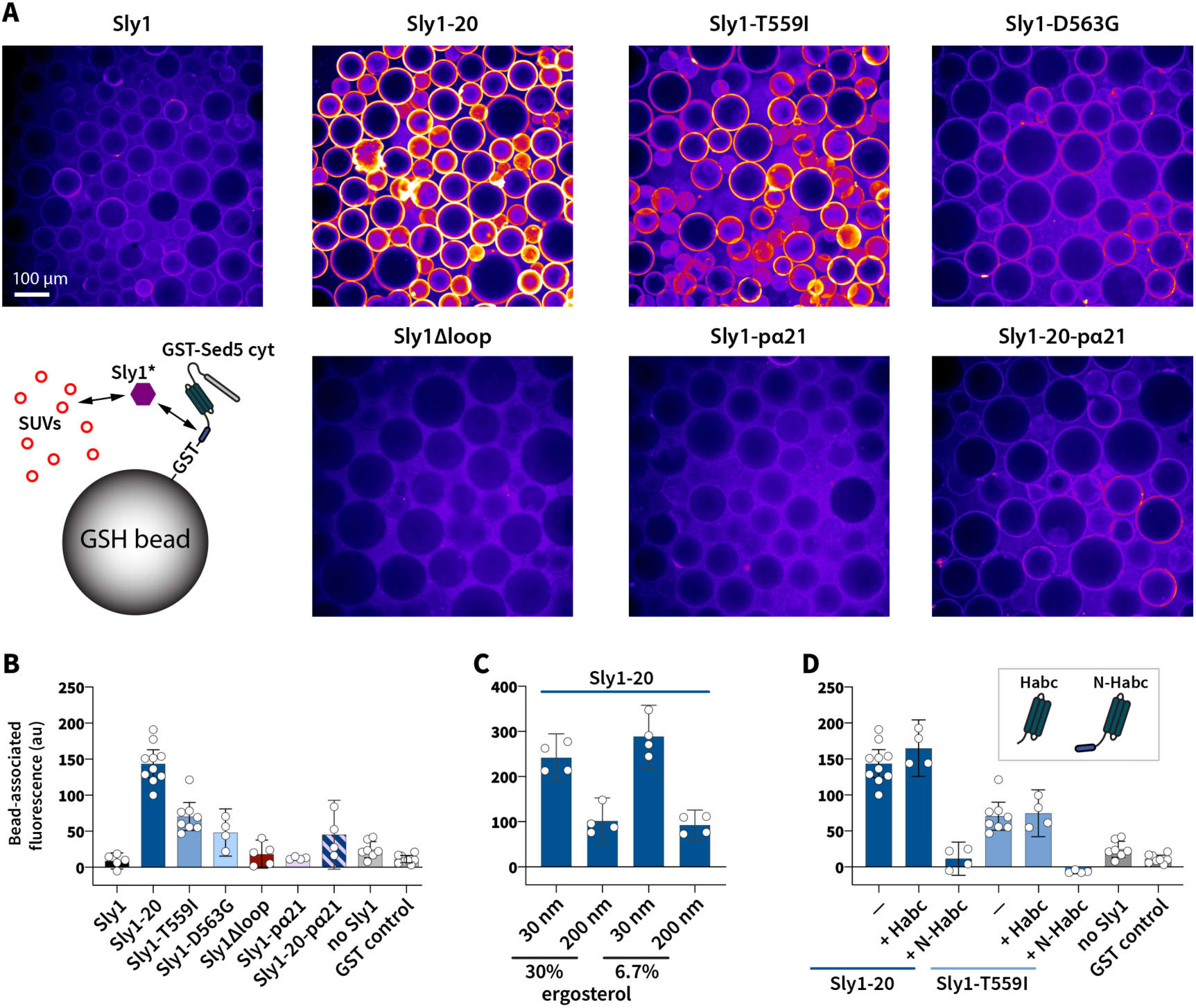
Hyperactive forms of Sly1 directly tether high-curvature vesicles to immobilized Sed5. **A**, The ability of Sed5-bound Sly1 to directly tether vesicles was tested using a bead-based assay system. GST-Sed5 was adsorbed to glutathione-Sepharose (GSH) beads, and wild-type or mutant forms of Sly1 were added to the reaction mixture. After 5 min, fluorescent R-SNARE RPLs bearing Sec22 were added to the mixtures, incubated for 15-20 min, and imaged by confocal microscopy (10× objective). A false-color scale was used to emphasize small differences in contrast under conditions with less tethering. The micrographs are representative of at least two independent assays per condition. **B-D**, To quantify tethering efficiency we used a spin-down assay. Binding reactions set up as for microscopy were subjected to low-speed centrifugation to sediment the glutathione-Sepharose (GSH beads) and associated proteins and vesicles. The supernatant was removed and detergent was added to the pellet to liberate bound fluorescent lipids; the resulting signal was quantified by fluorometry. In **C**, Sly1-20 was present for each condition. In **D**, Sly1* was pre-incubated with a 6:1 excess of soluble Sed5-Habc or Sed5-N-Habc, as indicated. Y-axes show bead-associated fluorescence (au, arbitrary units) after subtracting background from blanks containing only buffer. Bars indicate means ±95% confidence intervals for 4-10 independent experiments.

Sly1 binds the Sed5 N-terminal domain with sub-nM affinity (aa 1-21; Bracher and Weissenhorn, 2002; Demircioglu et al., 2014; Yamaguchi et al., 2002). To test the importance of this binding interaction in tethering, Sly1-20 was pre-incubated with a 6:1 molar excess of Sed5-N-Habc (aa 1-210; **Fig. 9D**). This abolished tethering. In contrast, tethering was not blocked by Sed5-Habc (aa 22-210), which lacks the N-peptide. Thus, to tether vesicles Sly1 must bind the N-peptide of the immobilized Qa-SNARE. Taken together the present and previously reported genetic data, and our assays of *in vitro* fusion, peptide binding, and tethering, all support the conclusion that the amphipathic helix α21 is necessary and sufficient for direct Sly1 binding to the incoming vesicle’s lipid bilayer. This binding both de-represses Sly1 and allows it to tether incoming vesicles at close range.

## DISCUSSION

*SLY1* was identified through isolation of the *SLY1-20* as a dominant single-copy suppressor of deficiency in *YPT1*, which encodes the yeast Rab1 ortholog. It soon became apparent that Ypt1 regulates ER-Golgi transport, and that *SLY1* gain-of-function alleles might become hyperactive through the loss of negative regulation. The present experiments strongly support that hypothesis, but further demonstrate that helix α21 has at least two functions. First, α21 is needed for relief of Sly1 autoinhibition. Second, α21 has a positive function. Both α21 functions are essential for bypass of tethering requirements by *SLY1-20*, and both require the presence of conserved apolar residues within α21. The same apolar residues are required for direct binding of α21 to membranes. Sly1 mutants lacking the loop, or with a constitutively open loop that has reduced membrane affinity, exhibit loss of function relative to the wild type. Thus, the loop’s ability to bind membranes has functions beyond relief of Sly1 autoinhibition.

In a working model (**Fig. 10A**), long-range tethers such as Uso1/p115 mediate initial capture of the vesicle, operating at ranges of 30-200 nm or more. Multisubunit Tethering Complexes (MTCs) including GARP, Dsl, and COG have long, floppy appendages (Chou et al., 2016; Ha et al., 2016; Ren et al., 2009). Golgins have an extended coiled-coil structure; their rod-like coiled-coil domains are interspersed with hinge-like domains. (Cheung and Pfeffer, 2016; Gillingham, 2018) In two cases, Golgin-210 and the Golgin-like endosomal tether EEA1, there is evidence that the hinges cause the tether to buckle or collapse, allowing the vesicle to approach the target membrane (Cheung et al., 2015; Murray et al., 2016). We propose that in the early secretory pathway long-range tethering factors hand vesicles off to Sly1, which would tether vesicles at a range of ∼15 nm from the target membrane to promote *trans*-SNARE complex assembly (**Fig. 10B**). The Sly1 loop’s preference for small-diameter vesicles in our binding and tethering assays is reminiscent of the behavior of ALPS domains, which seem to operate as selectivity filters that recognize bulk physical properties of membranes in the early secretory pathway (Bigay et al., 2005; Bigay and Antonny, 2012; Drin et al., 2008; Magdeleine et al., 2016). We propose that Sly1’s tethering function adds an additional layer of selectivity to this system.

**Fig. 10.**
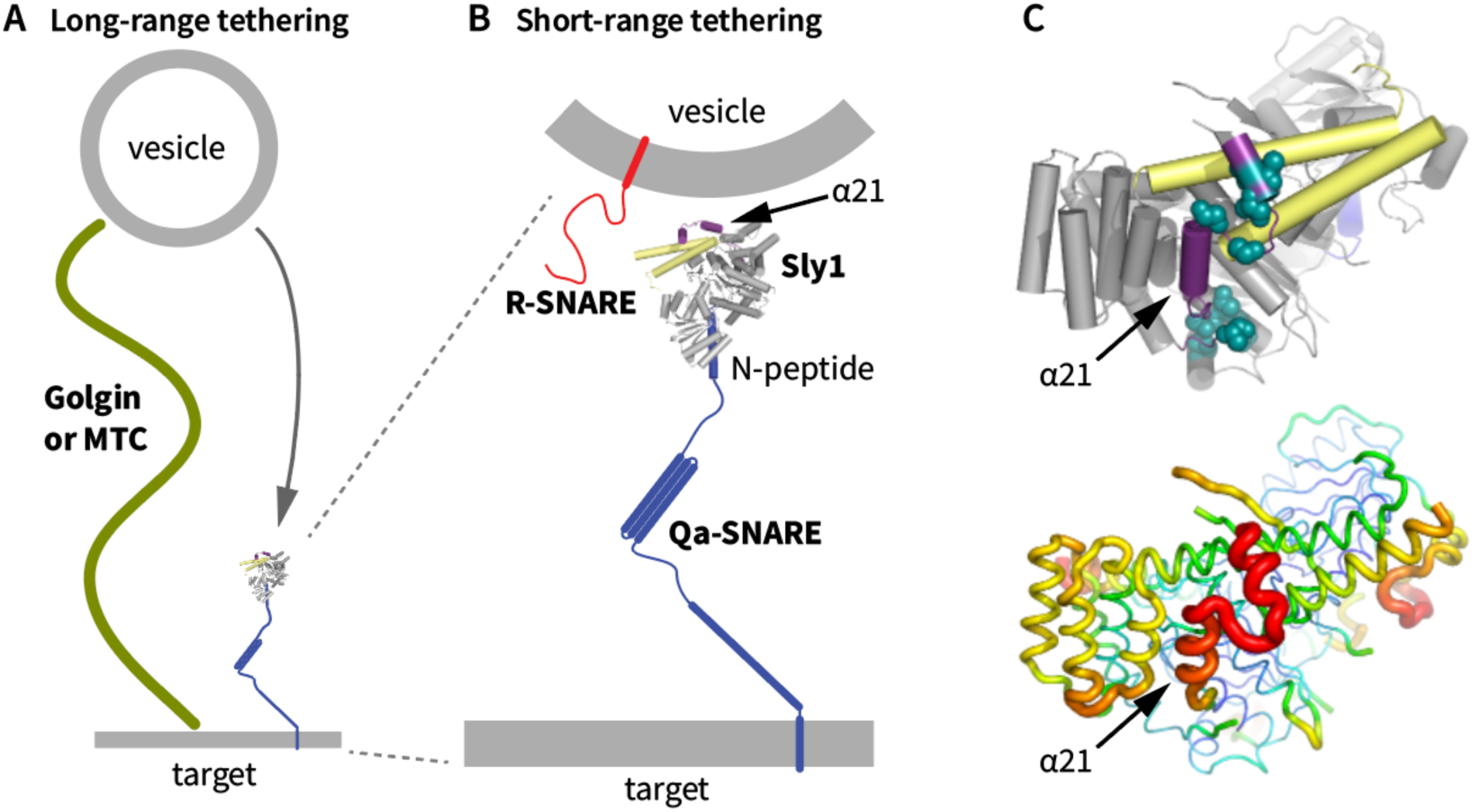
Working model. **A**, Long-range tethering is mediated by coiled-coil Golgin family tethers and multi-subunit tethering complexes (MTC’s). Flexibility or buckling of long-range tethers allows the vesicle to dwell in the region near Sly1 so that handoff can occur. **B**, Mechanism of close-range tethering. Sly1 is anchored to the N-terminal domain of the Qa-SNARE on the target membrane. Note that in the closed ground state, the loop and helix α21 sit opposite the N-peptide. Binding of α21 to an incoming vesicle’s membrane pulls open the autoinhibitory loop and tethers the vesicle to Sly1, likely in a spatial orientation optimal for R-SNARE binding to Sly1 domain 3a (yellow). **C.** The Sly1 loop is conformationally heterogenous in the crystal structure (PDB 1MQS). The top rendering shows locations of single gain-of-function amino acid substitutions Sly1, as in Fig. 1B. The bottom rendering shows relative temperature (B) factors, encoded by color and backbone trace thickness. The poorly conserved amino terminal half of the loop was not resolved in the deposited structure.

In the anterograde ER-Golgi pathway both Sly1 and the Qa-SNARE Sed5 must be present on the Golgi acceptor membrane; they cannot fulfill their functions if located only on COPII-derived transport vesicles (Cao and Barlowe, 2000). Sly1 is anchored to Sed5 through a direct, sub-nanomolar interaction with the Sed5 N-terminal domain (Bracher and Weissenhorn, 2002; Demircioglu et al., 2014; Yamaguchi et al., 2002). As our experiments show, Sly1 binding to the Sed5 N-peptide is indispensible for Sly1-mediated tethering (**Fig. 10B**). However, a previous report argued that the Sed5-Sly1 interaction is of relatively minor importance (Peng and Gallwitz, 2004). In the companion manuscript (Duan et al., submitted) we show, consistent with tethering experiments presented here, that deletion of the Sed5 N-peptide severely impairs fusion *in vitro* and is lethal *in vivo*.

Sitting almost exactly opposite Sly1’s N-peptide-binding cleft is the Sly1 loop (**Fig. 10B**). The loop is mobile: in the Sly1 crystal structure the poorly conserved N-terminal half of the loop is unresolved, while the better-conserved C-terminal half of the loop is partially resolved but exhibits a high temperature (B) factor (**Fig. 10C**), an indication of conformational heterogeneity. We speculate that when Sly1 is in its auto-inhibited ground state, helix α21 undergoes a “log-rolling” rotation about its long axis, intermittently exposing apolar side chains to probe for the presence of incoming vesicle membranes. Helix α21 binding to the vesicle bilayer has two consequences. First, the loop is pulled open, exposing the R-SNARE binding surface on Sly1. Second, the loop operates as a close-range tether, stabilizing the vesicle and target membrane within a distance sufficient to allow R-SNARE binding to Sly1 and the nucleation of a *trans*-SNARE complex on Sly1 domain 3a. The Sly1 loop might also constrain the rotational motion of Sly1 so that Sly1 is optimally oriented for productive R-SNARE binding. Although the available data are entirely consistent with this working model, we emphasize that many details are provisional and should be tested in future studies.

In our *in vitro* tethering assays, Sly1-20 and other gain-of-function mutants allow efficient tethering, consistent with the ability of these mutants to suppress requirements for other tethering factors. In the same *in vitro* assays, however, wild type Sly1 tethers much less efficiently. This raises the question of whether close-range tethering is important for wild type Sly1. The Sly1Δloop mutant cannot be auto-inhibited, yet it exhibits substantial tethering and fusion defects *in vitro*. *In vivo*, our SGA analyses revealed that the *sly1Δloop* allele exhibits synthetic sick or lethal interactions with dozens of genes involved in ER and Golgi traffic, including many genes that encode tethering factors or their regulators. In other words, when close-range tethering is prevented even partial defects in long-range tethering result in catastrophe and death. We suggest that a key function of Golgi long-range tethers is to allow incoming vesicles to dwell in the vicinity of Sly1 for long enough to allow inspection of vesicle membrane properties by α21, leading to loop opening, close-range tethering, R-SNARE binding, and assembly of a fusion-proficient *trans*-SNARE complex.

Additional mechanisms might contribute to the Sly1 loop’s function. First, it is possible that as-yet unidentified proteins bind Sly1, contributing to loop opening and tethering. Second, when open (as in Sly1-20), the loop may be intrinsically disordered, generating a “steric cushion” that locally exerts force on the adjacent docked membranes (Busch et al., 2015; D’Agostino et al., 2017). Our evidence for such a steric cushion mechanism is equivocal. *In vitro*, the behavior of Sly1Δloop, which completely lacks the loop, and of Sly1-20-pα20, which has a full-length loop that is constitutively open but partially defective in membrane binding, exhibit similar defects in most assays. This would argue against the steric cushion hypothesis. *In vivo*, however, *SLY1-20-pα20* allows almost wild type growth, while the *sly1Δloop* mutant grows slowly. Finally, it is possible that α21 binding to the vesicle locally perturbs its membrane structure, lowering the energy barrier for the onset of lipid mixing. Additional work will be needed to evaluate these potential mechanisms.

Sly1 has been proposed to promote vesicle fusion in several ways. (*i*) The Golgi Qa/t-SNARE Sed5 can adopt a tightly closed, autoinhibited conformation. Sly1 can open closed Sed5, allowing SNARE complexes to form more readily, at least in aqueous solution (Demircioglu et al., 2014; Kosodo et al., 1998). (*ii*) As we have shown here, helix α21 binding to membranes both de-represses and directly promotes Sly1 activity through a mechanism involving close-range vesicle tethering. (*iii*) Sly1 has conserved structural features that in Munc18-1 and Vps33 have been shown to catalyze *trans*-SNARE complex assembly through a Qa–R-SNARE templating mechanism. (*iv*) We have shown (again in aqueous solution but corroborated by genetic experiments) that Sly1 can decrease the rate of SNARE complex disassembly by Sec17 and Sec18. In the accompanying manuscript (Duan et al., submitted) we argue that that each of these mechanisms contributes to Sly1 function and that all are required for full Sly1 activity. In particular, we show that the close-range tethering mechanism characterized here is central Sly1’s ability to selectively nucleate *trans*-versus *cis-*SNARE complexes.

The Sly1 loop is conserved among Sly1 homologs from yeast to human but is absent from representatives of the three other SM sub-families: Sec1/Munc18, Vps45, and Vps33. Why is the loop unique to Sly1? We suggest that Sly1 must function in a considerably broader variety of cellular and molecular contexts than other SM’s. For example, the endosomal SM Vps45 associates with a scaffold protein, Vac1 (in mammals, Rabenosyn-5). Vac1 binds both Rab5 and phosphatidylinositol-3-phosphate, and might mediate close-range tethering in a manner analogous to the Sly1 loop (Burd et al., 1997; Peterson et al., 1999; Rahajeng et al., 2010; Tall et al., 1999). Similarly, Sec1 physically and functionally interacts with the exocyst tethering complex, and Vps33 is stably associated with Vps-C tethering complexes including HOPS and CORVET, which subsume both tethering and SNARE assembly functions (Morgera et al., 2012; Rieder and Emr, 1997). In the case of exocyst, the tethering activity is subject to autoinhibition, which is apparently released by engagement of rho family GTPases (Rossi et al., 2020).

In an interesting parallel, an ALPS-like domain within the HOPS subunit Vps41 was proposed to select high-curvature endocytic vesicles for docking and fusion (Cabrera et al., 2010). We also note that Munc18-1, despite lacking the Sly1-specific regulatory loop, is reported to tether vesicles in a reaction that requires at least the Qa-SNARE N-peptide and the R-SNARE on the opposing membrane (Arnold et al., 2017; Tareste et al., 2008). It is not clear whether Munc18-1 mediated tethering entails a direct interaction between Munc18-1 and the vesicle bilayer. This parallel may suggest that close-range tethering is a subreaction common to SM function.

Which specific long-range tethers hand vesicles off to Sly1 for close-range tethering, docking, and fusion? Persuasive experiments show that Sly1 operates in concert with Ypt1 and Uso1 (yeast Rab1 and p115, respectively) on the anterograde ER-Golgi pathway (Cao and Barlowe, 2000). However, direct interactions between Sly1 and Ypt1 or Uso1 have not been detected. Binding interactions have been detected between human Sly1 and COG, and perhaps between yeast Sly1 and Dsl (Kraynack et al., 2005; Laufman et al., 2009). However, the positive suppressing interactions of *SLY1-20*, the negative synthetic sick or lethal interactions of *sly1Δloop*, and the known SNARE interactions of Sly1, all point to Sly1 operating as “receiving agent” for vesicular traffic into the ER, early Golgi compartments, and perhaps later Golgi compartments as well. Additionally, several ER and Golgi tethers are reported to bind directly to Sly1 client SNAREs. Thus Sly1 might accept cargo containers presented by COG, GARP, TRAPP, Dsl, and the various Golgins. A significant challenge for the future is to identify the combinations of long-range tethers and SNAREs that operate in concert with Sly1, and to delineate the mechanisms that coordinate handoffs from long-range tethers to the core fusion machinery.

## MATERIALS & METHODS

### Yeast strains and *SLY1* gain-of-function screen

We use standard *Saccharomyces* genetic nomenclature (Dunham et al., 2015). Dominant alleles, whether wild type or mutant, are named in uppercase type (e.g., *SLY1-20*); recessive alleles are named in lowercase (*e.g.*, *sly1Δloop*). Strains and plasmids used in this study are described in **Supplementary Table 4**. To obtain new suppressors of *uso1Δ*, a library of *SLY1\** mutant alleles was constructed using the GeneMorph II Random Mutagenesis Kit (Agilent #200550). The *SLY1* open reading frame was amplified using the “medium mutation rate” PCR protocol. Four mutagenic PCR pools were separately purified and cloned into a derivative of the yeast vector pRS415, which contained 431 bp of the *SLY1* promoter and 249 bp of the *SLY1* terminator, using traditional restriction–ligation methods. Aliquots of the pRS415::*SLY1* mutant library ligation products were transformed into TOP10F’chemically competent *E. coli* cells, and 10 individual clones were Sanger sequenced to assess cloning fidelity and mutation frequency. Each clone sequenced contained the *SLY1* open reading frame with 0-4 mutations, with about 50% of the clones containing mutations. After the *SLY1* mutant library pools were verified, aliquots of the mutant library ligation products were transformed into Bioline Alpha-Select Gold Efficiency Competent *E. coli* cells. Transformant colonies were scraped from the LB + Amp transformation plates (maintaining four separate mutant pools), and allowed to grow for about two doublings. Plasmid DNA was extracted and purified from each of the pooled cultures using Qiaquick columns. 1ug of plasmid DNA from each *SLY1* mutagenic pool was transformed into *S. cerevisiae* strain AMY2144 (CBY1297: *uso1Δ* pRS426::*USO1*). Transformant colonies were grown under selection for leucine auxotrophy, then replica plated to synthetic complete medium containing 5-FOA, and incubated for 2 days. Yeast colonies that grew on 5-FOA (thus “kicking out” the WT copy of *USO1*) where struck out on –LEU plates, and plasmid DNA was purified from ten or more clones from each pooled library, using the Smash and Grab procedure. Plasmids were Sanger sequenced. On the basis of these results, pRS415::*SLY1* single mutant alleles were constructed. Site-directed *SLY1\** mutants were constructed using PCR and Gibson assembly, and are described in **Supplementary Tables 1–3**. The second half of *sly1-pα21* gene and its derivative *SLY1-20-pα21* (see **Supplementary Table 2**) were ordered as a gBlock (IDT) and cloned into the BamHI and NcoI sites on the wild-type *SLY1* plasmid.

### SGA analysis

A query strain (AMY2443) was constructed in the Y9205 genetic background (Tong and Boone, 2005), with *sly1Δloop* and a linked nourseothricin (*NAT*) marker integrated through allelic replacement at the native *SLY1* locus. This query strain was crossed to the *MAT* **a** haploid deletion and DAmP libraries, where each individual genetic perturbation is marked with a *KAN* resistance marker (Breslow et al., 2008; Tong and Boone, 2005). Diploids were selected by robotic pinning (Singer RoToR) onto YPD + 100 mg/L clonNAT + 200 mg/L G418, then induced to sporulate by pinning to sporulation medium (20g/L agar, 10g/L potassium acetate, 1g/L yeast extract, 0.5g/L glucose, 0.1g/L amino acid supplement [2g histidine, 10g leucine, 2g lysine, 2g uracil]) and growth at room temperature for 5 days. Spores were subsequently pinned to haploid selection medium (SD -His/Arg/Lys + 50 mg/L canavanine + 50 mg/L thialysine) and *MAT* **a** meiotic progeny grown for 2 days at 25° C. This haploid selection step was repeated, and the resulting colonies imaged using a Phenobooth (Singer) imaging system. These colonies encompass all potential meiotic progeny and serve as the control strains for phenotypic normalization. Haploid double mutants carrying both the *KAN* deletion allele and the *sly1Δloop*::*NAT* allele were selected by pinning meiotic progeny to double selection medium (SD/MSG -His/Arg/Lys + 50mg/L canavanine + 50 mg/L thialysine +100 mg/L clonNAT + 200 mg/L G418). After 2 days of growth at 25° C, this selection step was repeated and duplicate plates incubated at either 30° C or 37° C. Plates were imaged using the Phenobooth system, and colony size differences calculated using PhenoSuite software and web app (https://singerinstruments.shinyapps.io/phenobooth/).

### Protein purification

Full-length SNARE proteins were produced as previously described (Furukawa and Mima, 2014) with modifications. *E. coli* Rosetta2 (DE3) pLysS cells (Novagen) harboring each of the SNARE expression plasmids with 3C protease-cleavable N-terminal tags (pET-41/GST-His_6_ for SEC22 and pET-30/His_6_ for SED5, BOS1 and BET1) were inoculated from a 1:1000 dilution of the starter culture grown in MDAG-135 medium (Studier, 2005) into 1 L of Terrific Broth supplemented with 100 µg/mL Kanamycin and 34 µg/mL Chloramphenicol and grown at 37°C, 275 rpm until OD600 reached ∼1. Cultures were then induced with 1 mM IPTG for 3 h at 37°C. Cultures were harvested at 5000 × g and cell pellets were snap frozen with liquid nitrogen. Each liter yielded ∼10 g of wet cells, which were stored at −70°C. For purification, the frozen pellets were warmed to −10° C and broken up into small pieces with a metal spatula, then resuspended at a ratio of 5 mL of buffer per g of cell paste in 1× SNARE buffer (20 mM Na·PO_4_, 500 mM NaCl, 10% (m/v) glycerol, 1 mM DTT, pH 7) supplemented with 30-40 mM imidazole, 0.25 mg/mL chicken egg lysozyme, 125 U benzonase per g of cells, and 1× Sigmafast Protease inhibitor cocktail. 4 mL (1/10 volume) of 1 M n-octyl-β-D-glucopyranoside in H_2_O (β-OG, Anatrace) was added to 100 mM final concentration; the suspension was rotated at room temperature for 25 min to allow detergent-aided enzymatic lysis. Lysates were clarified at 16,500 × g, 4° C for 10 min, transferred to clean centrifuge tubes and centrifuged again for 20 min. Clarified lysates were batch-bound with 2ml of Ni-Sepharose HP equilibrated in 1× SNARE buffer with β-OG for 30 minutes. SNARE-bound resin was washed in plastic disposable columns with 25 mL of SNARE buffer supplemented with β-OG and 60-100 mM imidazole. SNARE proteins were eluted with SNARE buffer supplemented with β-OG and 200-300 mM imidazole, and snap frozen in liquid nitrogen. Purified protein was quantified using by absorbance at 280 nm and purity was assessed with SDS-PAGE with Coomassie blue staining. Protein aliquots were stored at −70° C until reconstitution. We note that 3C protease caused substantial unintended cleavage of Bos1 in its N-terminal linker domain due to a cryptic 3C site (148-GLPLYQ/GL-155). Mutation of the poorly-conserved residue Q153 to aspartic acid eliminated unintended proteolysis.

Soluble domains of Sed5 were expressed from the pET-30 vector (for H_6_-Habc and H_6_-N21-Habc) or pET-49 vector (for GST-H_6_-SED5ΔTM) and purified in same way as the full length protein except that the temperature was lowered to 35°C prior to induction, the buffers did not contain β-OG, and lysis was performed using Emulsiflex-C5 high pressure homogenizer (Avestin). Eluted protein was exchanged into FB160M1 (20 mM HEPES·KOH, 160 mM KOAc, 10% (m/v) Glycerol, 1mM MgOAc_2_, pH 7) using a PD-10 desalting column. Precipitated material was removed by centrifugation at 10,000 × g for 10 minutes and soluble protein aliquots were snap frozen in liquid nitrogen in 250ul PCR tubes and stored at −80° C. Sec22(SNARE)-GFP-His_8_ was expressed from the pST50Trc1 vector in Rosetta2(DE3) cells grown in ZYM-5052 autoinduction media (Studier, 2005) supplemented with carbenicillin (100 µg/mL) and chloramphenicol (34 µg/mL) overnight (>16 h) at 30° C from a 1:1000 dilution of starter culture. Cells were harvested and protein was purified as for soluble domains of Sed5. Sec17 was purified as described (Schwartz and Merz, 2009) except that the culture was grown in ZYM-5052 autoinduction media (Studier, 2005) at 37°C until OD_600nm_ was ∼0.8; temperature was then lowered to 18°C and the culture was incubated for ∼24 hours. Sec18 was purified as described (Haas and Wickner, 1996).

Sly1 and its mutants were expressed in Rosetta2(DE3) cells from pHIS-Parallel1 vectors (Lobingier et al., 2014; Sheffield et al., 1999). Frozen glycerol stocks were used to inoculate overnight starter cultures at 37° C in MDAG-135 containing 100 mg/L carbenicillin and 50 mg/L chloramphenicol (Studier, 2005). Each starter culture was diluted 1/1000 to seed 1-2 L of Terrific Broth containing 100 mg/L carbenicillin and 34 mg/L chloramphenicol. These cultures were grown in an orbital shaker (37° C, 275 rpm) to OD_600nm_ ∼1. Cultures were then transferred to a prechilled shaker at 16°C for 1 h before induction with 0.1-1 mM IPTG for 18 h. Cells were sedimented and resuspended in cold Sly1 buffer (20 mM Na·PO_4_, 500 mM NaCl, 10% (m/v) glycerol, 1 mM DTT, pH 7) supplemented with 30 mM imidazole, 0.25 mg/mL chicken egg lysozyme and 1× Sigmafast Protease inhibitor cocktail at a ratio of 5 mL of buffer per g of cell paste. The cells were lysed by passing through Emulsiflex-C5 high pressure homogenizer (Avestin) 2-4 times and the lysate was clarified by centrifugation at (16,500 × g, 25 min, 4° C). Clarified lysate from 1 L of culture (2 L for Sly1Δloop and Sly1-20) was bound in batch with 1 mL equilibrated Ni^2+^-Sepharose HP resin (GE) for 30 min at 4° C. Sly1-bound resin was collected in a 25 mL disposable Econo-Pac column (Bio-Rad) by gravity and washed with 25 mL of SLY1 buffer supplemented with 50 mM imidazole at pH 7. Sly1 was eluted with Sly1 buffer supplemented with 300 mM imidazole pH 7 in 0.5 mL fractions. Most of the protein eluted in fractions 3-7. Sly1 was exchanged into FB160M1 (20 mM HEPES·KOH, 160 mM KOAc, 10% m/v Glycerol, 1 mM MgOAc_2_, pH 7) using a PD-10 column (GE Healthcare). Precipitated material was removed by centrifugation at 10,000 × g for 10 minutes and soluble protein were diluted or concentrated to ∼2.4 mg/mL. Aliquots were snap-frozen in liquid nitrogen in thin-wall PCR tubes and stored at −70° C.

Recombinant HRV3C protease was prepared either as an N-terminal His8-tag fusion (AMP2019) or an N-terminal GST-His_6_-(Thrombin) fusion (AMP2016). 1 L of 1/1000 dilution of an overnight culture of Rosetta2(DE3) cells harboring the expression plasmid was grown overnight at 37° C in ZYM-5052 autoinduction media with 100 µg/mL kanamycin and 34 µg/mL chloramphenicol. Cells were centrifuged, resuspended in 4 times the volume of Lysis buffer (50 mM Tris-HCl pH 8.0, 300 mM NaCl, 10% glycerol, 1 mM DTT, and no protease inhibitors) supplemented with 15 mM imidazole and 0.5 mg/mL lysozyme, and lysed using Emulsiflex-C5 high pressure homogenizer (Avestin). Clarified lysate was incubated with 3 mL Ni^2+^-Sepharose HP (GE) for ∼30 min and strained in a disposable column. Resin was washed thoroughly with Lysis buffer supplemented with 40-60 mM imidazole and protein was eluted with 200 mM imidazole in about 7.5 mL. Concentrated fractions were combined and EDTA was added to 1 mM. The yield was ∼100 mg of purified protease per 1 L of culture. Purified protease was diluted to 10 mg/ml and exchanged into storage buffer (50 mM Tris-HCl (pH 8.0), 150 mM NaCl, 1 mM EDTA, 1 mM DTT, and 20% glycerol.), frozen in liquid N_2_, and stored at −80°C. Protease activity of the preparations was assayed using a homemade assay based on a linked FRET pair of fluorescent proteins (Evers et al., 2006), modified with an HRV 3C-cleavable linker. Reduction in FRET due to proteolysis was monitored in real time using a SpectraMax Gemini microplate reader (Molecular Devices).

GST-His_6_ was expressed and purified using conventional Ni^2+^ IMAC chromatography methods. Protein was exchanged into FB160M1 before freezing in liquid N_2_ and stored at −80° C.

### Circular Dichroism Spectroscopy

Purified SLY1wt or SLY1Δloop was exchanged into CD buffer (20 mM xNa-Pi, 100 mM NaCl, pH 7.2), diluted to 0.2 mg/mL, and loaded into a 0.1 cm path length cuvette. Spectroscopy was performed using a J-1500 CD Spectrophotometer (JASCO) at 25°C. CD and absorbance were measured from λ = 195 to 260 nm in steps of 0.1 nm. The protein concentration during each read was determined from absorbance at 205 nm, using the extinction coefficient at 205 nm calculated by the Anthis and Clore method (http://nickanthis.com/tools/a205.html) for each protein. Molar ellipticity for each protein was calculated by dividing the CD at each wavelength by the cuvette pathlength and protein concentration. Mean residue ellipticity for each protein was calculated by dividing the Molar ellipticity by the number of amino acids per protein.

### Preparation of RPLs

The FB160 buffer system and lipid mixtures used here are derived from B88 buffer, used extensively in COPII vesicle budding assays (Baker et al., 1988), and from lipidomic studies. The ER lipid mix is based on “Major-Minor” mixtures used for COPII budding (Antonny et al., 2001; Matsuoka et al., 1998), while the Golgi-mix is based on lipidomic surveys (Klemm et al., 2009; Schneiter et al., 1999). In particular, the study of Schneiter et al. used a highly enriched Golgi fraction known to be competent for docking and fusion of COPII carrier vesicles (Lupashin et al., 1996). Relatively high concentrations of ergosterol were used based prior work on COPII budding, which demonstrated that higher sterol levels yielded more morphologically homogenous COPII budding profiles (Matsuoka et al., 1998). In pilot studies, however, RPLs prepared with lower ergosterol concentrations exhibited similar fusion characteristics, including Sly1 and PEG dependence, as the high-sterol RPLs used in the experiments presented here. Lipids were obtained from Avanti Polar Lipids as chloroform stocks (Alabaster, AL) except for ergosterol, which was from Sigma-Aldrich. **Supplementary Table S5** lists the proportions, working stocks, and volumes of lipids and detergent used to prepare ER-mix and Golgi-mix RPLs. Lipid stocks were prepared or purchased in chloroform, except for ergosterol and phosphatidylinositol-4-phosphate, which were dissolved in 1:1 chloroform:methanol. β-OG stock solutions were prepared in methanol. Lipid-detergent films were prepared by transferring lipid and β-OG stocks to a glass vial (typically, 8 µmol total lipids and 70 µmol β-OG). The mixture was dried under a nitrogen stream; residual solvent was removed using a Speedvac™ evaporator. The lipid-detergent film was hydrated and solubilized with 400 µL 5× FB160M1 by three cycles of bath sonication and shaking. To the lipid– β-OG mixture, content mixing FRET reporters were then added (500 µL of 4 mg/mL solution of R-phycoerythrin-biotin conjugate, or 296 µL of 2mg/mL Alexa-Streptavidin; both reagents from Thermo/Molecular Probes). SNARE stocks in SNARE elution buffer with β-OG were then added to a final molar ratio of 1:600 (each Q-SNARE) or 1:300 (Sec22) to total phospholipids. Water was used to fill the headspace necessary to dilute 5× FB160M1 buffer to 1× concentration (2 mL final volume). Mixtures were nutated for 30 minutes before recombinant 3C protease was added (in 1:10 ratio to total SNAREs) to cleave affinity tags from the SNARE proteins during dialysis. The resulting mixtures were dialyzed (20 kDa cutoff) for ∼18h at 4°C in the dark against 250 volumes of FB160M1 containing 2 g BioBeads SM2 (Bio-Rad, Hercules, CA) per 2 mL of RPL mixture. The RPL mixture was then separated from unencapsulated content mixing probe by floating the RPLs up a step gradient of iso-osmotic Histodenz (35/25/0%) in FB160M1 (SW60Ti rotor at 55k rpm for 90 min), harvested, and diluted to 2 mM phospholipid. Phospholipid was quantified by measuring the fluorescence of the membrane fluorophore, initially verified by inorganic phosphate analysis (Chen et al., 1956). 32 µL aliquots of RPLs were transferred to thin-wall PCR tubes and frozen by immersion in liquid N_2_. RPLs prepared by this method and stored at −80° C were stable and fusion-competent, with minimal leakage of encapsulated FRET probes, for at least one year.

### RPL fusion assays

Unless noted otherwise, a standard order-of-addition was always used to initiate RPL assays. 250 µM (final phospholipid) of each RPL was premixed with PEG6K and other fusion components such as Sec17, Sec18 and ATP. Fusion assays were performed in 20 µL sample volumes in 384-well plates (Corning #4514). The reactions were monitored in a plate-based fluorimeter (Molecular Devices Gemini XPS or EM) for 5 min to establish a baseline, then Sly1 was added to initiate fusion. Except as noted, the moment of Sly1 addition was defined as time = 0. Lipid mixing was monitored with Ex_370nm_ and Em_465nm_. Content mixing was monitored with Ex_565nm_ and Em_670nm_. Graphs show mean ± s.e.m. of n ≥ 3 independent assays. Curves on the graphs show a second-order kinetic model fit to each dataset using a weighted least-squares algorithm in GraphPad Prism. Some experiments were run in both the presence and absence of unlabeled streptavidin, to assess leakage of RPL aqueous contents. For content mixing, typical signal for a complete reaction over background (*e.g.*, no Sly1) exceeded 50:1.

### Yeast growth assays

Yeast strains containing pRS426::*USO1* or pRS416::*YPT1* and pRS415::*SLY1* mutant plasmids were grown in –LEU liquid media, then diluted using a 48-pin manifold or a multichannel pipettor onto 5-FOA plates. The 5-FOA plates were grown at restrictive or non-restrictive temperatures, as indicated. Growth was scored relative to positive and negative control strains after 2-3 days.

### Peptide-liposome binding assay

To prepare small Texas RED-DHPE labeled unilamellar vesicles, lipid chloroform stocks were mixed using Hamilton syringes in glass vials, dried under a nitrogen stream, and residual solvent was removed in a Speedvac™ concentrator. The resulting lipid films were rehydrated with FB160M1 and either sonicated or extruded using an Avanti mini extruder with 0.03, 0.05 or 0.2 µm polycarbonate filters (Whatman). Peptides were custom-synthesized with a tetramethylrhodamine (TMR) fluorophore at the N-terminus, and were >98% pure by HPLC. The fluorophore is zwitterionic and does not change the net charge (+1) of the peptides. Emission spectra were acquired using a Molecular Devices Gemini XPS fluorescence spectrometer. FRET ratios were calculated as the ratio of fluorescence emission at 610 and 585 nm. The data were normalized by comparing each sample to the corresponding no-FRET condition (sum of the FRET signals for each peptide and SUV, acquired separately):

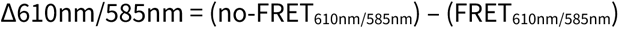

### Bead-based tethering assays

Beads were prepared in 100 µL (10 reactions) or 1 mL (100 reactions) batches. In small disposable spin columns, 100 µL of beads were washed in FB160M1 supplemented with 1% (m/v) bovine serum albumin (FB160M1BSA), and were loaded with 100 µg of GST-Sed5 cytoplasmic domain (1/5 of the resin’s nominal binding capacity; 150 pmol protein per 10 µL resin in a 1× reaction), in a volume of 500 µL FB160BSA; this mixture was incubated, with slow agitation, for 30 min at room temperature. Unbound material was removed by gentle centrifugation (∼70 × g, 10 seconds), the beads were washed once with FB160BSA, and the beads were then blocked by adding excess recombinant GST-His_6_ protein (1.25 mg; 2.5 × the resin’s nominal binding capacity) in FB160BSA, in 500ul final volume. Unbound GST-H6 was not removed. The bead-SED5-GST suspension was stored at 4° for up to a week. For tethering assays, 1× reaction aliquots of the bead-SED5-GST suspension (50 µL, containing ∼10 µL packed beads) were transferred to 250 µL PCR tubes, then Sly1* (75 pmol; a 1:2 molar ratio to Sed5_cyt_) was added to each reaction tube in 50 µL volume, allowing the Sly1* to bind to the immobilized GST-Sed5_cyt_ in 100 µL final volume. For competition experiments the Sly1* was pre-incubated with a 6-fold molar excess (450 pmol) of Sed5 Habc or N-Habc domain for 10 min at room temperature before adding the Sly1*-competitor mixture to the beads. Tethering was initiated by adding Texas-red-DHPE labeled SUVs to each 1× tethering reaction (1-6 µL depending on stock concentration). The tethering reactions were incubated for 15-20 min at room temperature, then transferred to wells of chambered coverslips that had been pre-incubated with FB160BSA for at least 20 min. These preparations were observed at ambient temperature (23±2° C) using a Nikon Ti2 microscope equipped with a Yokogawa CSU-X1 spinning disk confocal unit, a Toptica iChrome MLE laser combiner and launch; 405, 488, 561, and 647 nm diode lasers (Coherent); a Finger Lakes high-speed emission filter wheel; and a Mad City piezoelectric Z-stage. The microscope was controlled by Nikon Elements software and data analysis and figure preparation was done with the Fiji package of Image/J software and plug-ins. Tethering reactions were observed using a 10× 0.30 NA Plan Fluor objective and an Andor 888 EMCCD camera operated at an EM gain of 300 with 200 ms exposure per frame.

Tethering was also quantified using bead spin-down assays. Binding reactions were initiated as in the microscopy-based tethering experiments. To quantify SUVs tethered to the beads, the beads were washed once in 1.3 mL of FB160BSA and then sedimented for 1 min at 500 × g in a swinging-bucket rotor. The supernatant was carefully removed, and resin-bound lipids were eluted from the beads with 50 µL of BugBuster protein extraction reagent (Millipore). The beads were again sedimented. To quantify the amount of eluted TRPE lipid, 20 µL of the final supernatant was analyzed in a plate-reading fluorimeter (Molecular Devices Gemini XPS or Gemini EM; excitation 595 nm; cutoff 610 nm; emission 615 nm).

## ACKNOWLEDGEMENTS

We are grateful to Drs. M. Ailion, J. Bai, R. Baker, J. Cattin, S. Hoppins, I. Topalidou, and M. Zick for helpful advice and critical comments on the manuscript; C. Barlowe, C. Boone, and D. Waugh for antibodies and strains, D. Baker and the University of Washington Institute for Protein Design for computational resources, and D. Beacham (Molecular Probes/Thermo Fisher) for gifts of fluorescent reagents. These studies were supported by NIH/NIGMS R01 GM077349 and the University of Washington (AM), NIH/NIGMS T32 GM007270 (UN), NIH MARC T34 GM083883 (BD) and Medical Research Council MC_UP_1201/10 (EM).

## SUPPLEMENTARY MATERIAL

**Supplementary Fig. S1.**
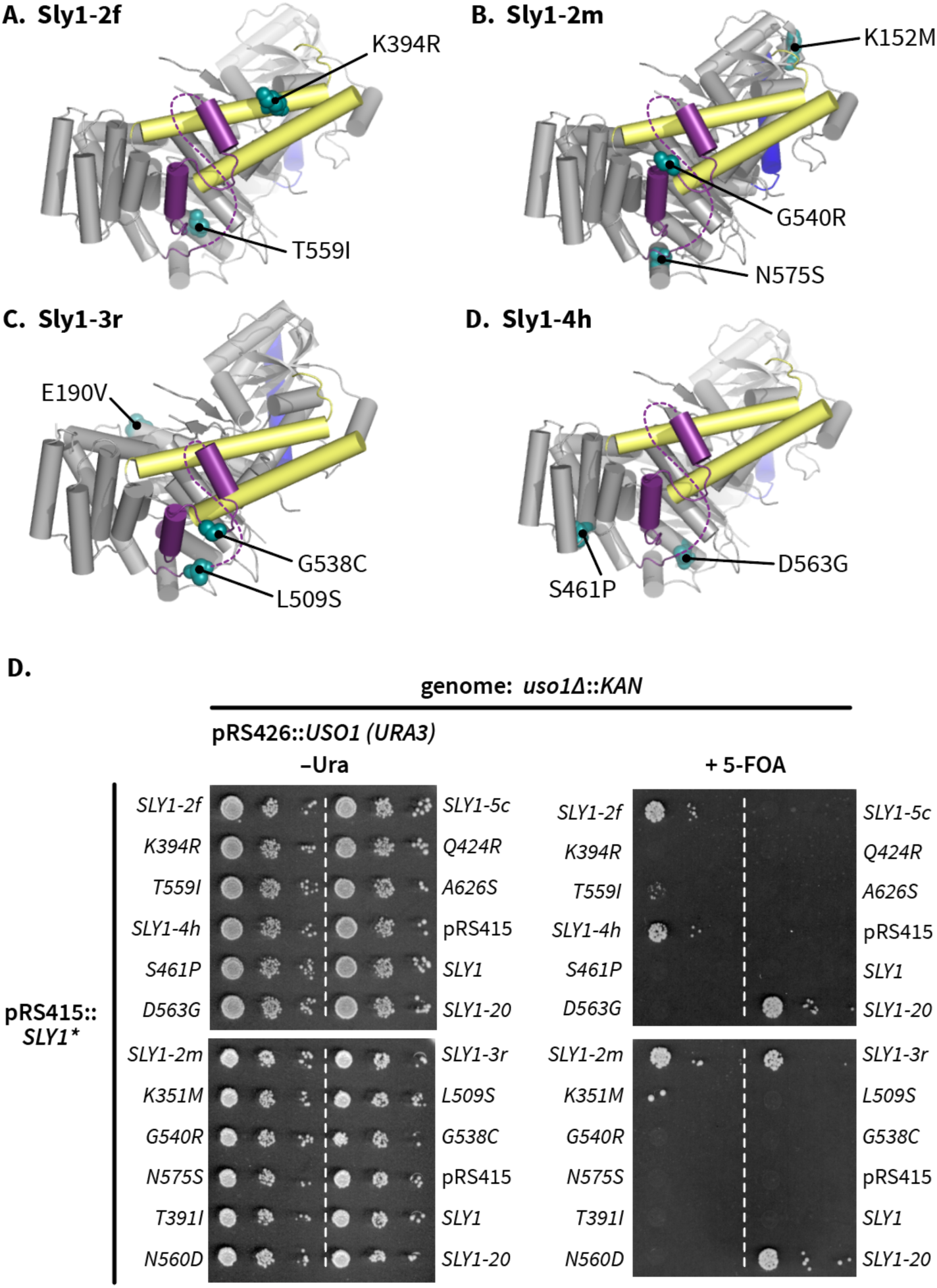
Some *SLY1* alleles require multiple substitutions to suppress the lethal *uso1*Δ phenotype. **A-D**, Locations of amino acid substitutions in four representative *SLY1* alleles recovered in our screen. **E**, Growth phenotypes show that most single substitutions are unable to suppress the total absence of Uso1. Many of the same single mutants suppress deficiency of Ypt1 (see Supplementary Table I). The multisite allele *SLY1-5c*, although retrieved in our primary screen, was unable to suppress the *uso1*Δ allele in secondary screening.

**Supplementary Table 1.**
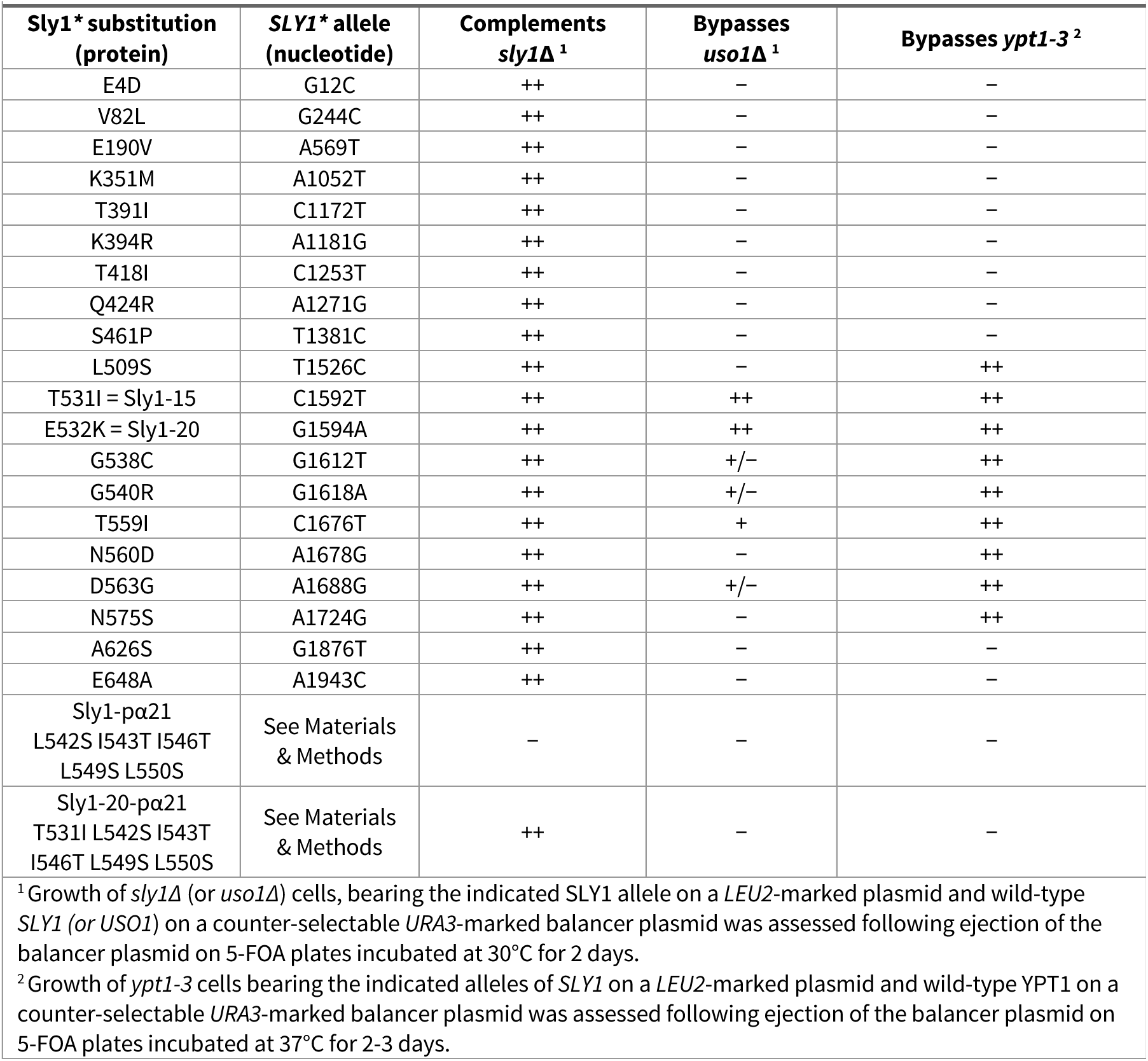
Selected *SLY1* mutants and their growth phenotypes.

**Supplementary Table 2.**
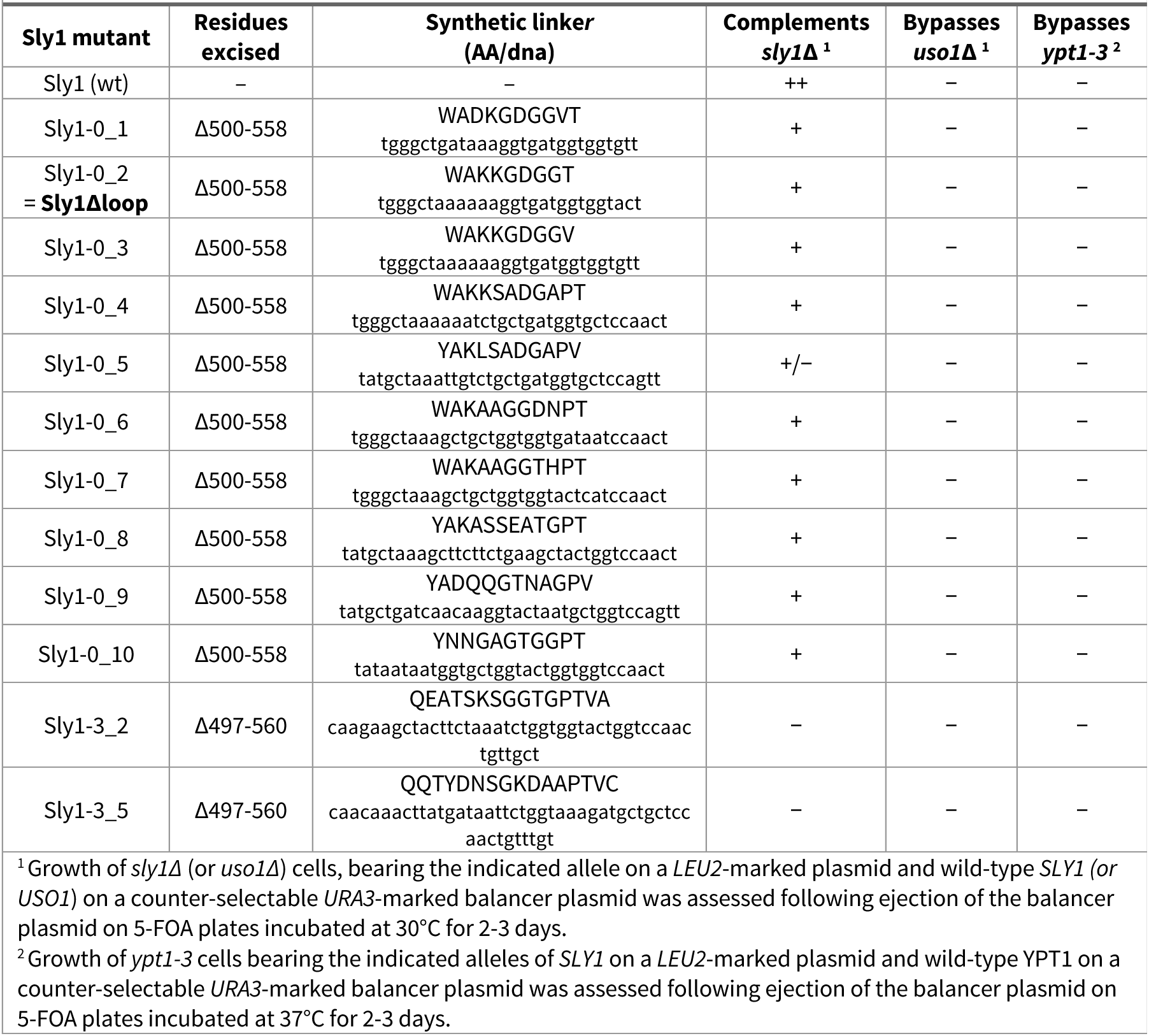
*SLY1* “loopless” mutants and their growth phenotypes.

**Supplementary Table 3.**
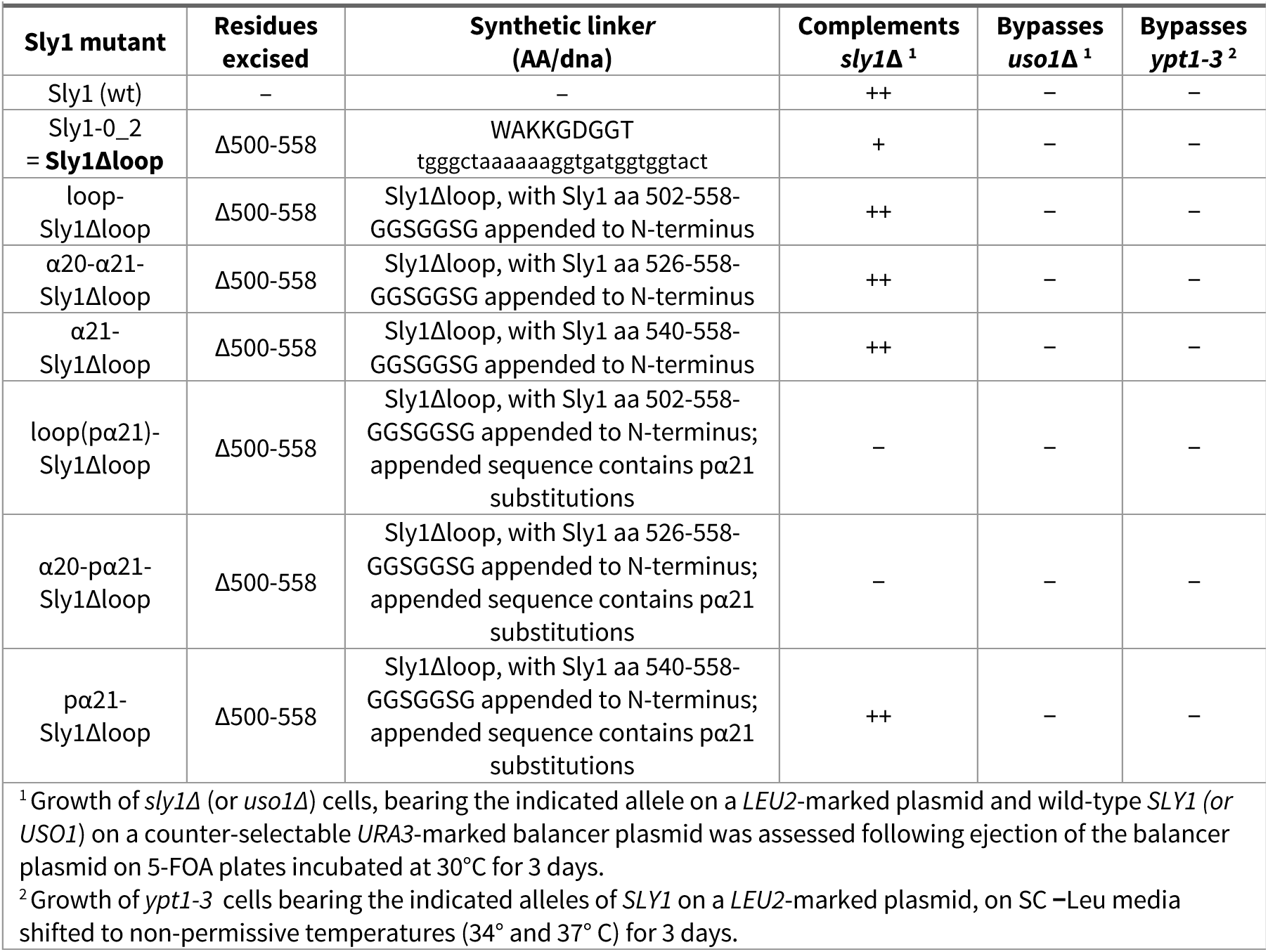
*SLY1* “loopless” chimeras with N-terminal Loop attachments, and their growth phenotypes.

**Supplementary Table 4.** Yeast strains and plasmids used in this study

| Strain/plasmid ID | Description/genotype | Source/reference |
| --- | --- | --- |
| **Yeast Strains** |  |  |
| AMY2143 (CBY481) | *MATalpha ura3-1 trp1-1 ade2-1 leu2-3,112 can1-100 ypt1-3* | C. Barlowe |
| AMY2349 | *MATalpha ura3-1 trp1-1 ade2-1 leu2-3,112 can1-100 ypt1-3 CBB1395 (URA3 2μm YPT1)* | C. Barlowe |
| AMY2144 (CBY1297) | *MATalpha ura3Δ his3Δ leu2Δ met15Δ lys2Δ uso1Δ::KAN pSK47 (URA3 2μm USO1)* | C. Barlowe |
| AMY2232 (CBY73) | *MATa ura3-52 his3Δ200 trp1Δ63 ade2-101 lys2-801 sly1Δ::HIS3 AMP1525 (URA3 CEN SLY1)* | C. Barlowe |
| AMY2141 (Y8205) | *MATalpha ura3Δ0 his3Δ1 leu2Δ0 can1Δ::STE2pr-Sp_his5 lyp1Δ:: STE3pr-LEU2* | C. Boone |
| AMY2440 | *MATalpha ura3Δ0 his3Δ1 leu2Δ0 can1Δ::STE2pr-Sp_his5 lyp1Δ:: STE3pr-LEU2 SLY1-20 (NatMX6)* | This study |
| AMY2441 | *MATalpha ura3Δ0 his3Δ1 leu2Δ0 can1Δ::STE2pr-Sp_his5 lyp1Δ:: STE3pr-LEU2 SLY1-T559I (NatMX6)* | This study |
| AMY2442 | *MATalpha ura3Δ0 his3Δ1 leu2Δ0 can1Δ::STE2pr-Sp_his5 lyp1Δ:: STE3pr-LEU2 SLY1-D563G (NatMX6)* | This study |
| AMY2443 | *MATalpha ura3Δ0 his3Δ1 leu2Δ0 can1Δ::STE2pr-Sp_his5 lyp1Δ:: STE3pr-LEU2 sly1Δloop (NatMX6)* | This study |
| AMY2580 | *MATalpha ura3Δ0 his3Δ1 leu2Δ0 can1Δ::STE2pr-Sp_his5 lyp1Δ:: STE3pr-LEU2 SLY1 WT (NatMX6)* | This study |
| **Yeast expression plasmids** |  |  |
| AMP1910 | pRS415::SLY1 | This study |
| AMP1578 | pRS415::SLY1-20 | This study |
| AMP1588 | pRS415::SLY1(T559I) | This study |
| AMP1589 | pRS415::SLY1(D563G) | This study |
| AMP2052 | pRS415::sly1Δloop | This study |
| AMP1911 | pRS415:: sly1-pa21 | This study |
| AMP1912 | pRS415:: sly1-20-pa21 | This study |
| Various | pRS415::SLY1<sup>*1</sup> | This study |
| pSK47 | pRS426::USO1 | C. Barlowe |
| CBB1395 | pRS426::YPT1 | C. Barlowe |
| pDN524 | pRS424 with additional restriction sites flanking 2μ origin | Lobingier et al., 2014 |
| pDN317 | pDN524::SEC17 | Lobingier et al., 2014 |
| pDN367 | pDN524::sec17-FSMS (F21S, M22S) | This study |
| **E. coli SNARE expression plasmids** |  |  |
| AMP1792 | pET-30::His<sub>6</sub>-(3C)-Sed5 | (Furukawa and Mima, 2014) |
| AMP1961 | pET-30:: His<sub>6</sub>-(3C)-Sed5Habc(22-210) | This study |
| AMP1960 | pET-30:: His<sub>6</sub>-(3C)-Sed5N-Habc(1-210) | This study |
| AMP1973 | pET-49::GST- His<sub>6</sub>-(3C)-Sed5cyt(1-319) | This study |
| AMP2020 | pET-30:: His<sub>6</sub>-(3C)-Bos1(Q153D) | (Furukawa and Mima, 2014) but modified |
| AMP1794 | pET-30:: His<sub>6</sub>-(3C)-Bet1 (with corrected missense mutation) | (Furukawa and Mima, 2014) but corrected |
| AMP1795 | pET-41::GST- His<sub>6</sub>-(3C)-Sec22 | (Furukawa and Mima, 2014) |
| **E. coli SNARE chaperone expression plasmids** |  |  |
| AMP1547 | pTYB12::intein-CBD-Sec17 | (Schwartz and Merz, 2009) |
| AMP77 | pQE9:: His<sub>6</sub>-SEC18 | (Haas and Wickner 1996) |

| ***E. coli* Sly1 expression plasmids** |  |  |
| --- | --- | --- |
| AMP1649 (pBL51) | pHIS-Parallel1:: His<sub>6</sub>-(TEV)-Sly1 | (Lobingier and Merz, 2014) |
| AMP1651 | pHIS-Parallel1:: His<sub>6</sub>-(TEV)-Sly1-20 | This study |
| AMP1652 | pHIS-Parallel1:: His<sub>6</sub>-(TEV)-Sly1(T559I) | This study |
| AMP1653 | pHIS-Parallel1:: His<sub>6</sub>-(TEV)-Sly1(D563G) | This study |
| AMP1654 | pHIS-Parallel1:: His<sub>6</sub>-(TEV)-Sly1Δloop | This study |
| AMP1936 | pHIS-Parallel1:: His<sub>6</sub>-(TEV)-α20-α21-Sly1Δloop | This study |
| AMP1937 | pHIS-Parallel1:: His<sub>6</sub>-(TEV)-α21-Sly1Δloop | This study |
| AMP1939 | pHIS-Parallel1:: His<sub>6</sub>-(TEV)-α20-pα21-Sly1Δloop | This study |
| AMP1940 | pHIS-Parallel1:: His<sub>6</sub>-(TEV)-pα21-Sly1Δloop | This study |
| AMP1932 | pHIS-Parallel1:: His<sub>6</sub>-(TEV)-Sly1-pα21 | This study |
| AMP1933 | pHIS-Parallel1:: His<sub>6</sub>-(TEV)-Sly1-20-pα21 | This study |
| ***E. coli* miscellaneous expression plasmids** |  |  |
| AMP1881 | pET-49::GST- His<sub>6</sub>-(3C) | This study |
| AMP2019 | pET-30:: His<sub>8</sub>-HRV3C(Protease) | This study |
| AMP2016 | pET-49::GST- His<sub>6</sub>-(Thrombin)-HRV3C(Protease) | This study |
| AMP203 | MBP-(TEV)- His<sub>6</sub>-TEV | (Kapust et al., 2002) |
| AMP2018 | pET-28- His<sub>6</sub>-CFP(3C)YFP (for HRV3C protease FRET assay) | This study |
| <sup>1</sup>applies to all SLY1 mutants described in supplementary tables S1, S2 and S3 |  |  |

**Supplementary Table 5.** SNARE RPL lipid compositions used in this study

| **Supplementary Table 5. SNARE RPL lipid compositions used in this study** |  |  |  |  |
| --- | --- | --- | --- | --- |
| **ER mimetic RPLs (R-SNARE)** |  |  | **Golgi mimetic RPLs (Qabc-SNARE)** |  |
| **Lipid** | **mol%** |  | **Lipid** | **mol%** |
| 18:1 (Δ9-Cis) PC | 35.0% |  | 18:1 (Δ9-Cis) PC | 35.0% |
| 16:0-18:1 PE | 16.0% |  | 16:0-18:1 PE | 14.0% |
| Soy PI | 6.3% |  | Soy PI | 6.3% |
| 16:0-18:1 PS | 5.6% |  | 16:0-18:1 PS | 5.6% |
| 16:0-18:1 PA | 3.5% |  | 16:0-18:1 PA | 3.5% |
| 18:1 CDP DG | 1.4% |  | 18:1 DG | 1.4% |
| Brain PI(4)P | 2.0% |  | Brain PI(4)P | 2.0% |
| Ergosterol | 30.0% |  | Ergosterol | 30.0% |
| 16:0 Marina Blue™ DHPE | 0.2% |  | 16:0 NBD-PE | 2.2% |

